# RTP801 REGULATES MOTOR CORTEX SYNAPTIC TRANSMISSION AND LEARNING

**DOI:** 10.1101/2020.10.15.340851

**Authors:** L Pérez-Sisqués, N Martín-Flores, M Masana, J Solana, A Llobet, J Romaní-Aumedes, M Canal, G Campoy, E. García-García, N Sánchez-Fernández, S Fernández-García, JP Gilbert, MJ Rodríguez, H-Y Man, E Feinstein, D Williamson, D Soto, X Gasull, J Alberch, C Malagelada

**Affiliations:** Department of Biomedicine, Faculty of Medicine, University of Barcelona, Catalonia, Spain; Institut de Neurociències, University of Barcelona, 08036 Catalonia, Spain; IDIBAPS- Institut d’Investigacions BiomèdiquesAugust Pi i Sunyer, Barcelona, 08036 Catalonia, Spain; Centro de Investigación Biomédica en Red sobre Enfermedades Neurodegenerativas (CIBERNED), Barcelona, 08036 Catalonia, Spain; Department of Biology, Pharmacology and Experimental Therapeutics, Boston University, Boston, MA, 02215USA; Quark Pharmaceuticals, Fremont, CA, 94555 USA; Kinesiology Program, School of Behavioral Sciences and Education, Penn State Harrisburg, Middletown, PA,17057 USA

**Keywords:** GluA1, motor learning, mTOR, Plasticity, RTP801

## Abstract

RTP801/REDD1 is a stress-regulated protein whose upregulation is necessary and sufficient to trigger neuronal death in *in vitro* and *in vivo* models of Parkinson’s and Huntington’s diseases and is up regulated in compromised neurons in human postmortem brains of both neurodegenerative disorders. Indeed, in both Parkinson’s and Huntington’s disease mouse models, RTP801 knockdown alleviates motor-learning deficits.

Here, we investigated the physiological role of RTP801 in neuronal plasticity. RTP801 is found in rat, mouse and human synapses. The absence of RTP801 enhanced excitatory synaptic transmission in both neuronal cultures and brain slices from RTP801 knock-out (KO) mice. Indeed, RTP801 KO mice showed improved motor learning, which correlated with lower spine density but increased basal filopodia and mushroom spines in the motor cortex layer V. This paralleled with higher levels of synaptosomal GluA1 and TrkB receptors in homogenates derived from KO mice motor cortex, proteins that are associated with synaptic strengthening. Altogether, these results indicate that RTP801 has an important role modulating neuronal plasticity in motor learning.

## INTRODUCTION

Synaptic plasticity is the ability to fine tune neuronal connectivity and dynamics upon demand, for example in situations in which individuals have to adjust movements in challenging environments. This process is known as motor learning and involves the acquisition of a novel motor skill that, once learned, persists after training period ends (Peters *et al*, 2017; Sanes & Donoghue, 2000; Xu *et al*, 2009).

The central hub for motor learning is the motor cortex, an interconnected structure with other brain regions such as the striatum, the thalamus, brainstem or the spinal cord (reviewed in (Shepherd, 2013; Shepherd & Huganir, 2007)). The complex process of acquiring new motor skills induces synaptic plasticity in the motor cortex and requires dendritic spine formation, consolidation and/or elimination, all leading to a necessary synaptic remodeling and strengthening (Peters *et al*, 2017; Sanes & Donoghue, 2000; Fu *et al*, 2012; Xu *et al*, 2009). Pyramidal neurons from the motor cortex and striatal medium spiny neurons (MSNs) predominantly undergo plastic changes along motor learning (Costa *et al*, 2004; Tjia *et al*, 2017). Regarding the motor cortex, projection pyramidal neurons from Layer V (LV) are the main excitatory input to the striatum involved in the corticostriatal pathway (Costa *et al*, 2004; Shepherd & Huganir, 2007; Hintiryan *et al*, 2016; Anderson *et al*, 2010). These plastic changes leading to motor learning involve, at least, increased levels of α-amino-3-hydroxy-5-methyl-4-isoxazolepropionic acid receptors (AMPAR) at dendritic spines (Kida *et al*, 2016; Roth *et al*, 2020) . However, the mechanisms by which these events are regulated are not yet clearly elucidated.

In many neurodegenerative diseases, along with neurological and psychiatric symptoms, motor dysfunction is a hallmark of disease progression. Among these disorders, we find Parkinson’s disease (PD), Huntington’s disease (HD), or amyotrophic lateral sclerosis, (Shepherd, 2013). Motor dysfunction is due, in part, to an impairment in the synaptic plasticity of the circuitries that control movement by interconnecting motor cortex and basal ganglia and the thalamus, and also the cerebellum (Guo *et al*, 2015; Xu *et al*, 2017; Calabresi *et al*, 2007, 2000).

RTP801/REDD1, coded by the *DDIT4* gene, is a stress-regulated protein that is sufficient and necessary to induce neuron death (Shoshani *et al*, 2002; Malagelada *et al*, 2006). It is elevated in cellular and animal models of PD in response to dopaminergic neurotoxins (Malagelada *et al*, 2006; Ryu *et al*, 2005) and is highly up regulated in neuromelanin positive neurons in the substantia nigra pars compacta (SNpc) of both sporadic (Malagelada *et al*, 2006) and parkin mutant PD patients (Romani-Aumedes *et al*, 2014). RTP801 induces neuron death by a sequential inactivation of mTOR and the survival kinase Akt (Malagelada *et al*, 2008) via the tuberous sclerosis complex 1/2 (TSC1/2). Regarding HD, RTP801 levels are highly increased in HD human brains, in differentiated neurons derived from induced pluripotent stem cells (iPSC) from HD patients (Martín-Flores *et al*, 2016) and in striatal synapses from HD mouse models (Martín-Flores *et al*, 2020). Besides, in neuronal models of the disease, RTP801 mediates mutant huntingtin (mhtt)-induced toxicity (Martin-Flores *et al*, 2015). Importantly, RTP801 contributes to motor-learning dysfunction in HD since RTP801 knockdown prevents from the appearance of motor learning deficits in the R6/1 model of the disease (Martín-Flores *et al*, 2020). This suggests that synaptic RTP801 deregulation is a common hallmark in neurodegeneration. Indeed, RTP801 coding gene *DDIT4* was recently described as one of the top three common deregulated transcripts in *postmortem* brain samples from PD and HD patients (Labadorf *et al*, 2018). Furthermore, RTP801 is sufficient to cause neuronal atrophy and depressive-like behavior (Ota *et al*, 2014) and it has a regulatory role in cortical development, neuronal differentiation (Malagelada *et al*, 2011) and peripheral nervous system myelination (Noseda *et al*, 2013). However, its physiological role in synaptic plasticity has not been resolved yet. For this reason, here we investigated the potential synaptic function of RTP801 in the corticostriatal pathway. By using cellular and murine models and *postmortem* human brains and performing behavioral, histological, electrophysiological and biochemical analysis, our results describe the implication of RTP801 in motor learning plasticity.

## RESULTS

### RTP801 is localized in the synapses of murine and human samples and modulates synaptic transmission *in vitro*

We first explored whether RTP801 was localized in synapses and whether it was involved in synaptic function, connectivity and transmission. Hence, we first isolated cortical and striatal crude synaptosomes from adult *postmortem* human brain, adult rat and mouse brains and from cultured rat cortical neurons. In all samples we observed the presence of RTP801 or its enrichment in crude isolated synaptic terminals in comparison to the initial homogenates (**Fig 1 A**), corroborating our own previous results (Martín-Flores *et al*, 2020). Interestingly, in cultured cortical neurons, we observed that RTP801 was expressed in the soma, dendrites and dendritic spines (**Fig 1 B**).

**Figure 1.**
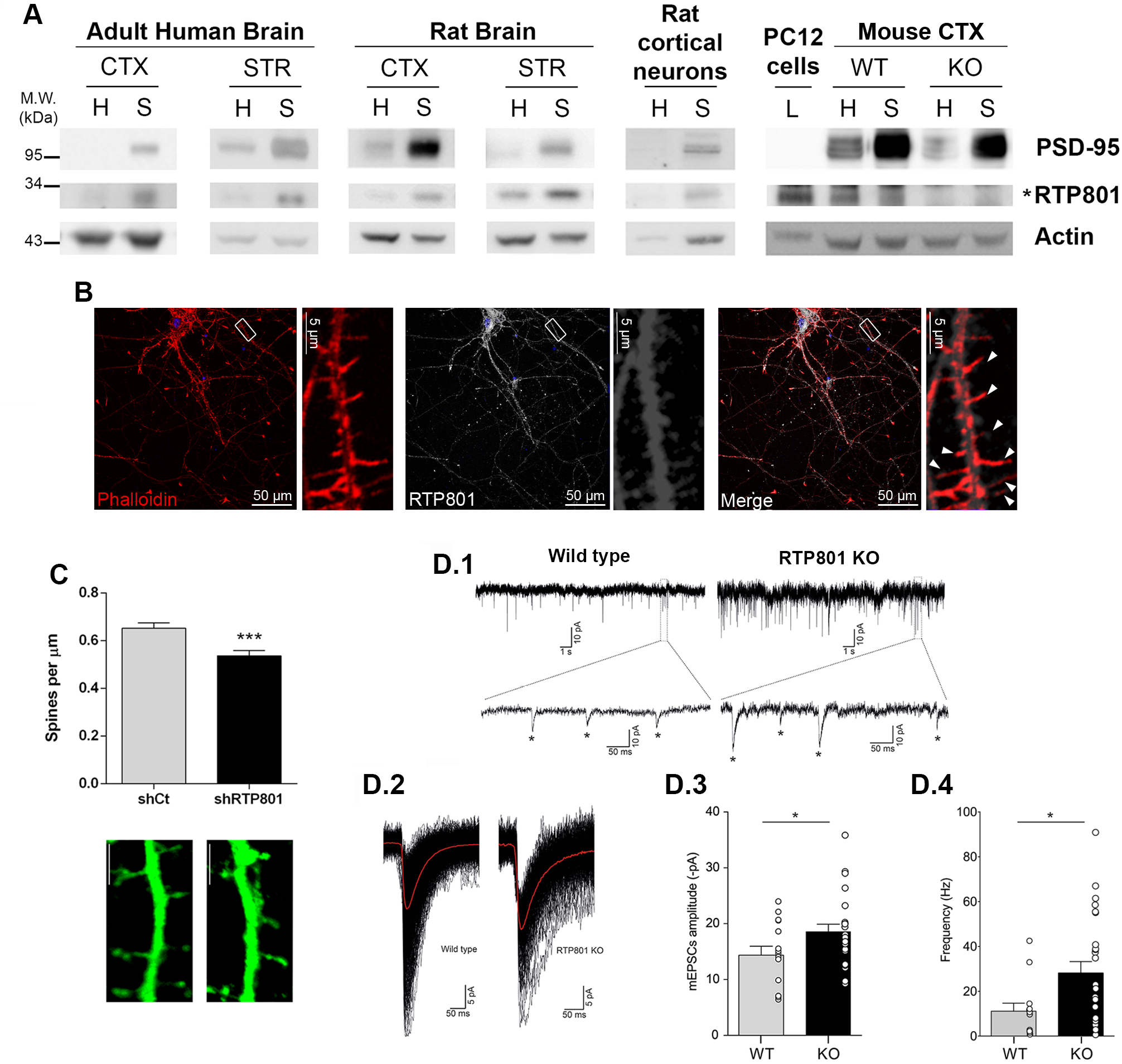
RTP801 is present at the synapse and modulates neuronal transmission. A. RTP801 protein is found in the synaptic compartment. Homogenate (H) and crude synaptic fraction (S) were obtained from human *postmortem cortex* (*CTX*) *and putamen* (*STR*), adult rat cortex (CTX) and striatum (STR), primary cortical cultures at 14 DIVs and cortex from both 2-months old WT and RTP801 KO mice. Whole cell lysate (L) from NGF-differentiated PC12 cells was added as a positive control to detect RTP801 in mouse brains. Samples were analyzed by western blot and proved against RTP801 (specific band pointed out by *), postsynaptic protein PSD-95 and actin as a loading control. **B. RTP801 is present ubiquitously in neurons, including at the synapses.** Primary rat cortical cultures were fixed at 14DIVs and stained for RTP801 (grey). Phalloidin (red) was used to visualize actin cytoskeleton. Nuclei were stained with Hoechst33342 (in blue). White arrows point F-actin-labeled dendritic spines colocalizing with endogenous RTP801 staining. **C. RTP801 knockdown reduces spine density in cultured cortical neurons.** Primary rat cortical neurons were transduced with lentiviral particles carrying a GFP-tagged control shRNA or an shRNA against RTP801. 4 days later (14 DIVs), cells were fixed and analyzed by immunofluorescence against GFP (green). *Scale bar, 5μm*. **D. Abrogation of RTP801 expression modulates synaptic plasticity *in vitro*. D.1.** Representative 20 seconds whole-cell recording of mEPSCs at a membrane voltage of −70mV from WT or RTP801 KO mice cultured cortical pyramidal neurons (14 DIV). A magnification (0.5 seconds) for both traces is shown below where asterisks denote the detected events. **D.2.** Example of averaged mEPSCs (red lines) superimposed on the individual mEPSCs (in black) from a wild type (average of 962 events) and RTP801 knockout (average of 236 events) culture. **D.3.** RTP801 KO recordings show differences in mEPSCs mean amplitude. **D.4.** The frequency of detected events in RTP801 KO neurons was statistically increased compared with WT. mEPSCs frequencies were obtained from same recordings shown in D.3. All data is presented as mean ± SEM from the recordings performed in 15 WT neurons and 25 KO neurons from at least six independent neuronal cultures. Statistical analyses for spine density and mEPSC amplitude were performed with Student’s t-test, *P< 0.05, ***P0.001 vs. shCt/WT and with Mann-Whitney test for mEPSC frequency, *P<0.05 vs. WT.

We next investigated whether RTP801 depletion affected spine density and synaptic transmission. For this, we knocked down the expression of RTP801 in cortical primary cultures at 14DIV, using lentivirus expressing a specific shRNA for RTP801 or scramble shRNA as control. We observed that RTP801 silencing induced a significant decrease in spine density relative to the scramble shRNA transduced neurons (**Fig 1 C**). We next analyzed whether RTP801 expression abrogation affected synapse function by evaluating the frequency and the amplitude of mEPSCs of cortical cultures derived from WT and RTP801 KO mice. Interestingly, in the complete absence of RTP801 expression using cultured cortical neurons from RTP801 KO mice, we observed that both the amplitude (**Fig 1 D1, D2 & D.3**) and frequency (**Fig 1 D1. D2 & D.4**) of mEPSCs were higher than the ones registered in WT cortical sister cultures.

We corroborated our *in vitro* results using cultured hippocampal neurons, a well characterized plasticity model. In line with previous results, we found that RTP801 colocalized with PSD-95, an excitatory postsynaptic scaffold protein, but not with the presynaptic marker SV2A, indicating that RTP801 is localized in the postsynaptic compartment (**Fig S1 A-B**). Moreover, ectopic RTP801 expression attenuated the amplitude of mEPSCs without affecting the frequency, along with a decrease of PSD-95 and AMPAR receptor subunit GluA1 puncta intensity (**Fig S1 C-E**).

### Synaptic and behavioral characterization of RTP801 KO mice brains

Previous data pointed out that the total abrogation of RTP801 expression did not influence significantly either the brain structure or the basal behavior of the RTP801 KO mice in comparison to WT animals (Brafman *et al*, 2004; Ota *et al*, 2014). However, we previously demonstrated that RTP801 regulated the timing of cortical neurogenesis and neuron differentiation/migration (Malagelada *et al*, 2011) using *in utero* electroporation techniques. For this reason, to validate the use of the RTP801 KO mouse to study its putative synaptic role, we characterized its brain morphology in comparison to WT animals. We first confirmed the lack of RTP801 expression in the KO animals in motor cortex homogenates (**Fig 2 A**). Macroscopically, although there were no differences in the mice body weight between genotypes (**Fig 2 B**), we observed that KO animals presented a decreased brain weight (**Fig 2 C**). However, internal structural organization did not present major alterations either in cortical layers, hippocampus or even in the striatum, as judged by Nissl staining (**Fig 2 D**). Primary motor cortex (M1) layer thickness did not differ either between genotypes (**Fig 2 E**) but RTP801 KO mice showed an expected decreased cell density in the M1 LV (**Fig 2 F**).

**Figure 2.**
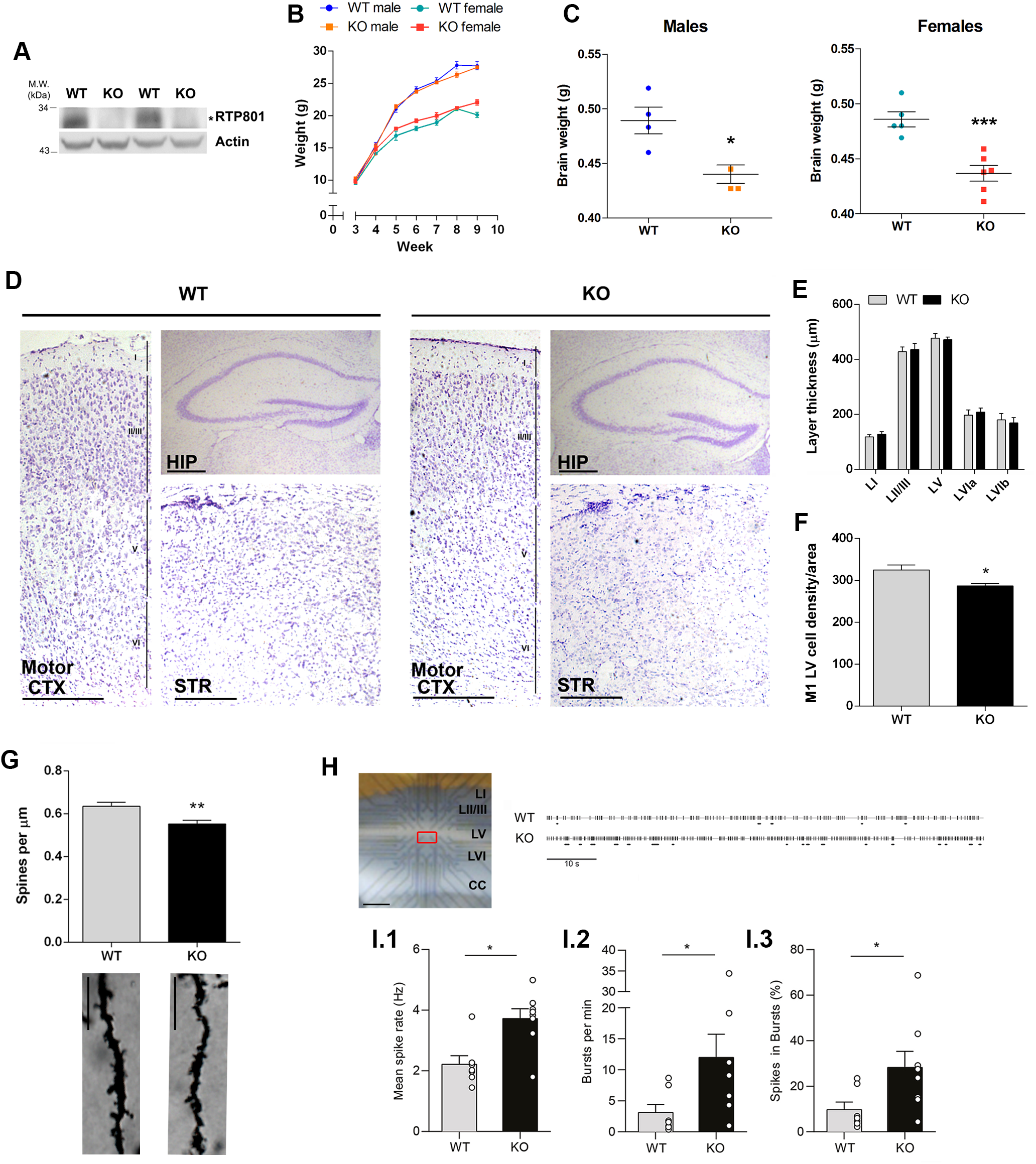
RTP801 KO mice show decreased brain size, decreased cortical spine density and enhanced synaptic transmission. **A.** Whole brain extracts from WT and RTP801 KO mice at 2 months of age were subjected to western blot. Membranes were proved against RTP801 (specific band is pointed out by *) and actin as loading control. **B.** 9-weeks long follow-up shows that RTP801 KO animals display normal weight. **C.** RTP801 KO mice exhibit decreased brain weight. Whole brain weight was measured in 2-months-old animals. **D**. Brain sections of WT and KO mice were subjected to Nissl staining to visualize cell somas. Representative images of motor cortex (CTX), hippocampus (HIP) and striatum (STR) for both genotypes are shown. *Scale bar*, 250μm. Motor cortex thickness (**E**) and LV cell density (**F**) were quantified in Nissl-stained sections. **G**. RTP801 total knockout mice show decreased spine density in motor cortex layer V. Spine density was quantified in 30 dendrites/animal, 50% apical and 50% basal, in 5 WT and 5 KO animals. *Scale bar*, 10 μm. **H.** Image of a brain sagittal slice on the MEA (magnification). Recordings from the selected electrodes (in red), located on LV, were analyzed. Motor cortex layers I-VI and *corpus callosum* (CC) are indicated. *Scale bar*, 600 μm. **I.** Spontaneous activity by MEA: Illustrative long time-scale (90 min) spike rasters of recorded LV motor cortex spontaneous activity from WT (4 males and 3 females) and RTP801 KO (4 males and 4 females) mouse brain slices (1 slice per animal); the horizontal lines above each raster define bursts. Graphs show quantification of spike rate (**I.1**), burst rate (**I.2**), and the percent of spikes that form bursts (**I.3**) of field spontaneous activity recorded by MEA. Data in all graphs are presented as mean ± SEM. * *P*<0.05, ***P<0.001; two-tailed Student t-test *versus* WT (**C**, **F**). *P<0.05; Mann-Whitney test versus WT (**I.1-I.3**). Data in (**B, E**) was analyzed by two-way ANOVA.

We next investigated whether cortical spine density was affected in the adult brain of RTP801 KO mice using Golgi-Cox staining. Analyses were performed in the M1 layer V pyramidal neurons, the main excitatory and direct projection to the ipsi- and contralateral striatum in the corticostriatal pathway (Shepherd, 2013; Hintiryan *et al*, 2016; Anderson *et al*, 2010; Xu *et al*, 2009). As previously seen by knocking down RTP801 in cortical cultured neurons (see **Fig 1 C**), we observed a reduction in the density of spines in LV neurons in naive RTP801 KO mice compared to WT animals (**Fig 2 G**).

Next, we assessed whether RTP801 modulate synaptic transmission in cortical brain slices from naïve WT and KO animals. We thus measured neuronal spike rate and bursting in M1 LV using multielectrode array (MEA) (**Fig 2 H**). We found an increased spike rate in the LV of KO animals when compared with WT (**Fig 2 I.1**), with no differences between male and female animals (**Fig S2 A**). Analysis of spike-train patterns showed a higher burst rate and proportion of spikes included in bursts in KO primary motor cortex slices when compared with WT (**Figure 2 I.2-I.3**). We found no other differences in the burst parameters analyzed (**Fig S2 B-D**). These results support the hypothesis that neuronal excitability is increased in LV motor cortex in KO mice as an attempt to compensate the decreased number of synaptic spines.

To study whether synaptic structural and functional changes in RTP801 KO mice correlated with behavioral alterations, we next investigated whether the lack of RTP801 affected coordination, locomotion and motor learning. We first tested WT and KO mice for hindlimb clasping, a marker of disease progression in a number of mouse models of neurodegeneration, including HD (Mangiarini *et al*, 1996; Chou *et al*, 2008). We observed that RTP801 KO male mice displayed a clasping phenotype, not present in male WT mice. The tendency in females was similar but not significant (**Fig S3 A**). We next explored whether gait, as a measure of coordination and muscle function, was affected in RTP801 KO mice. These animals showed a decrease in the length of the stride, stance, sway and the overlap (**Fig 3 A-B** and **Fig S3 B**), suggesting gait impairment in the KO animals. We next examined whether general locomotor activity was altered using the Open Field test. Despite gait impairment, we did not find any differences in the total distance travelled in the RTP801 KO mice relative to WT (**Fig 3 C**). We did not find differences in the distance travelled in the center or the time spent in the center, suggesting that RTP801 KO mice do not exhibit anxiety-like behavior. Regarding other general exploratory and stereotypic behavior, we did not find any differences in grooming or wall and vertical rearing, either (**Fig S3 C**).

**Figure 3.**
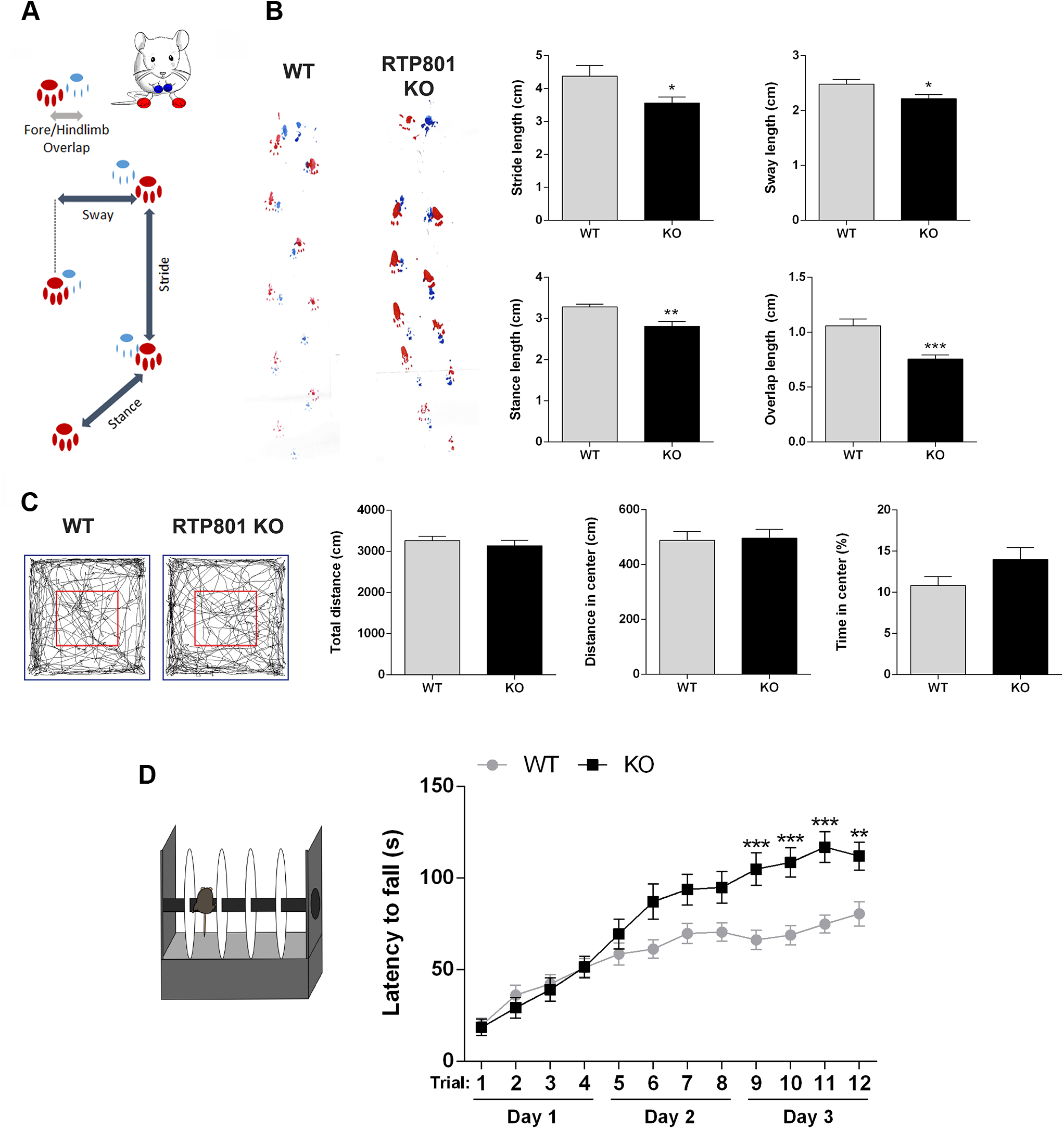
RTP801 contributes to motor learning and gait but does not alter general locomotor activity. **A.** Schematic representation of the four different parameters measured in the footprint test: stride, sway, stance and limbs overlap (in blue, forelimb prints and, in red, hindlimb prints). **B.** Representative examples of footprint tracking from both genotypes. Graphs on the right show hindlimb lengths for stride, sway, stance and limbs overlap. Data is represented as mean ± SEM and was analyzed with two-tailed Student’s t-test. * *P*< 0.05, ** *P*< 0.01, *** *P*< 0.001 *versus* WT group. N = 13 WT (8 males + 5 females) and 12 KO (4 males + 8 females). **C.** Representative tracking of mice activity recorded for 10 minutes in an open field test. Graphs on the right show total distance traveled in the whole arena (blue), distance traveled in the center (red) and percentage of time spent in the center. Measures are shown as mean ± SEM. There are no statistically significant differences according to the Student’s t-tests performed. N = 18 WT (6 males + 12 females) and 17 KO (7 males + 10 females). **D**. WT and RTP801 KO mice were subjected to the accelerating rotarod test and the time spent on it was evaluated for three days, four trials per day. Data is represented as mean ± SEM and was analyzed by two-way ANOVA followed by Bonferroni’s multiple comparisons test for *post hoc* analyses. Genotype effect: ** *P*< 0.01. Multiple comparisons: * *P*< 0.05, ** *P*< 0.01, *** *P*< 0.001 *versus* WT group in each trial. N = 31 WT (14 males + 17 females) and 30 KO (12 males + 18 females).

To evaluate motor skill learning, we trained the WT and RTP801 KO animals in the accelerating rotarod. Both female and male KO mice showed the same trend to improve motor learning in this behavioral paradigm (**Fig S3 D**). Together, RTP801 KO mice significantly improved performance in this task compared to WT animals (Genotype effect, ** *P*=0.0058) (**Fig 3 D**). This result indicates that RTP801 is involved in motor learning acquisition.

### RTP801 modulates spine density and structure in the primary motor cortex of trained animals

We next investigated whether the improvement in motor learning in the RTP801 KO mice affected differentially spine density and structure. Hence, since motor learning plasticity involves projections from the motor cortex to the dorsal striatum, we explored spine density and morphology in pyramidal neurons from the M1 LV and in medium spiny neurons (MSNs) from the dorsal striatum, one week after finishing the accelerating rotarod test (**Fig 4 A**). Similar to non-trained naïve RTP801 KO mice, trained RTP801 KO mice showed a decrease in the density of spines in LV pyramidal neurons (**Fig 4 B**), specifically in their basal dendrites. Interestingly, spine density of either cortical LV apical dendrites or dendrites in striatal MSNs did not change (**Fig 4 C-D**).

**Figure 4.**
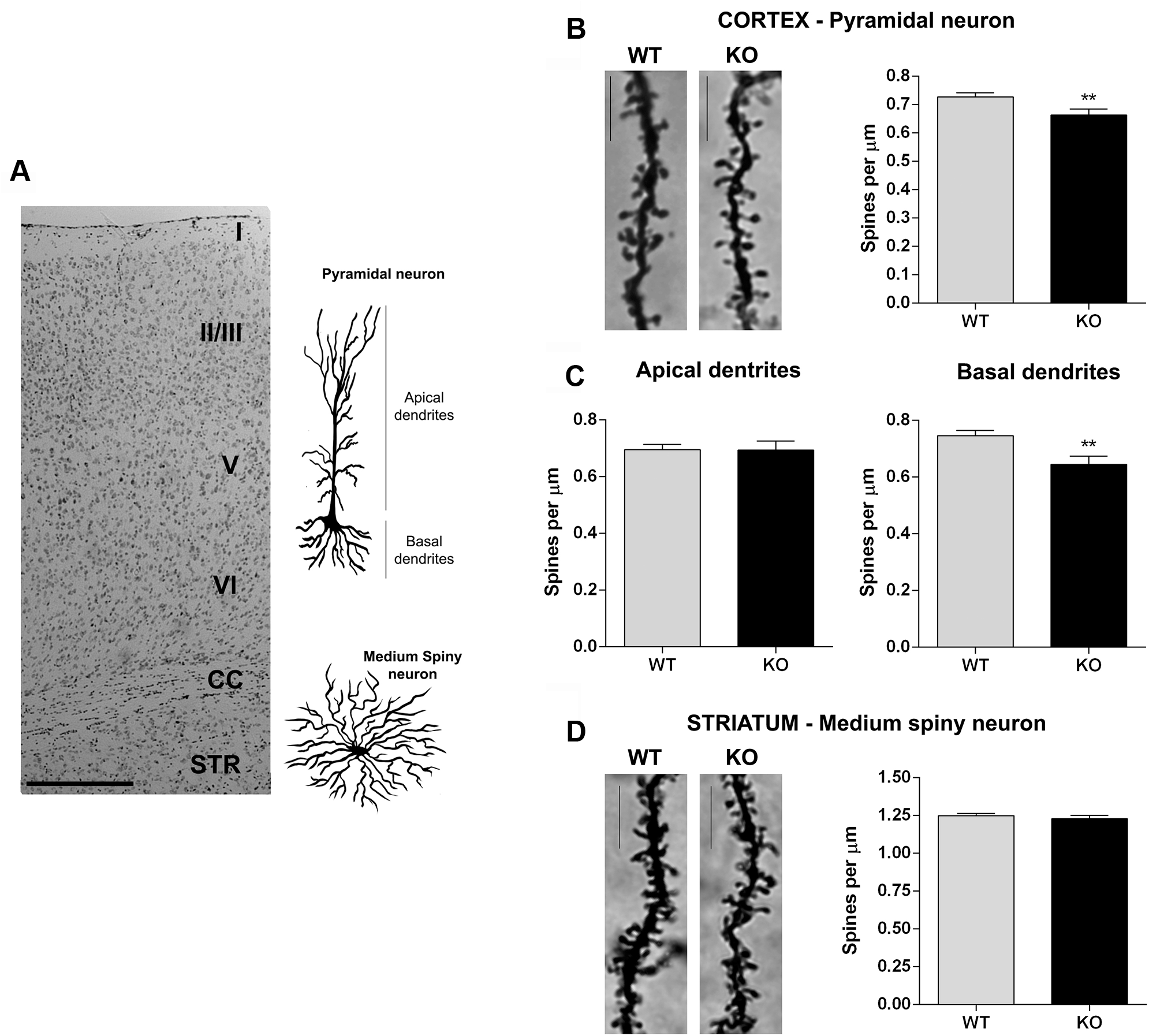
RTP801 modulates dendritic spine density and morphology of pyramidal neurons in M1 in motor-trained animals. One week after the accelerating rotarod test, WT and RTP801 KO mice were culled and their brains were impregnated with Golgi-Cox staining prior to analyze changes in spine density. **A, B. Loss of RTP801 expression decreases spine density of pyramidal neurons from M1.** Spine density was quantified combining both apical and basal dendrites from M1 LV pyramidal neurons. **A**, **C. Abrogation of RTP801 expression affects specifically basal dendrites of pyramidal neurons in M1.** Spine density was assessed in apical and basal dendrites. Each bar of the graphs represents mean ± SEM of at least 30 dendrites per animal (N= 8 WT and 5 KO), approximately 50% apical and 50% basal. **A, D. Loss of RTP801 expression does not affect spine density in the striatum.** Spine density was also analyzed in striatal MSNs. Data in the graph represent mean ± SEM of at least 30 dendrites per animal (N= 8 WT and 5 KO). Statistical analyses in a were performed with Mann-Whitney test, \*\**P*<0.01 vs. WT, and by Two-tailed Student’s t-test in c-d; ** *P*< 0.01 *versus* WT. Cortical layers (I, II/III, V, VI), *corpus callosum* (CC) and striatum (STR) are depicted in **A**. Representative WT and KO dendrites from primary motor cortex and striatum are shown (**B, D**). *Scale bar*, 10 μm.

Based on these results, we investigated differences in spine morphology in the M1 LV pyramidal neurons that could explain the increased motor learning in the KO mice. Indeed, RTP801 KO animals displayed more filopodia but less branched spines (**Fig 5 A** and **Fig S4**). In line with this, when related with the total number of headed spines, we observed a higher percentage of mushroom spines in the basal dendrites of KO animals (**Fig 5 B.1, C.1**). Moreover, their head area was also increased. On the contrary, no differences were found in either the percentage or head area of thin spines (**Fig 5 C.1, C.2**).

**Figure 5.**
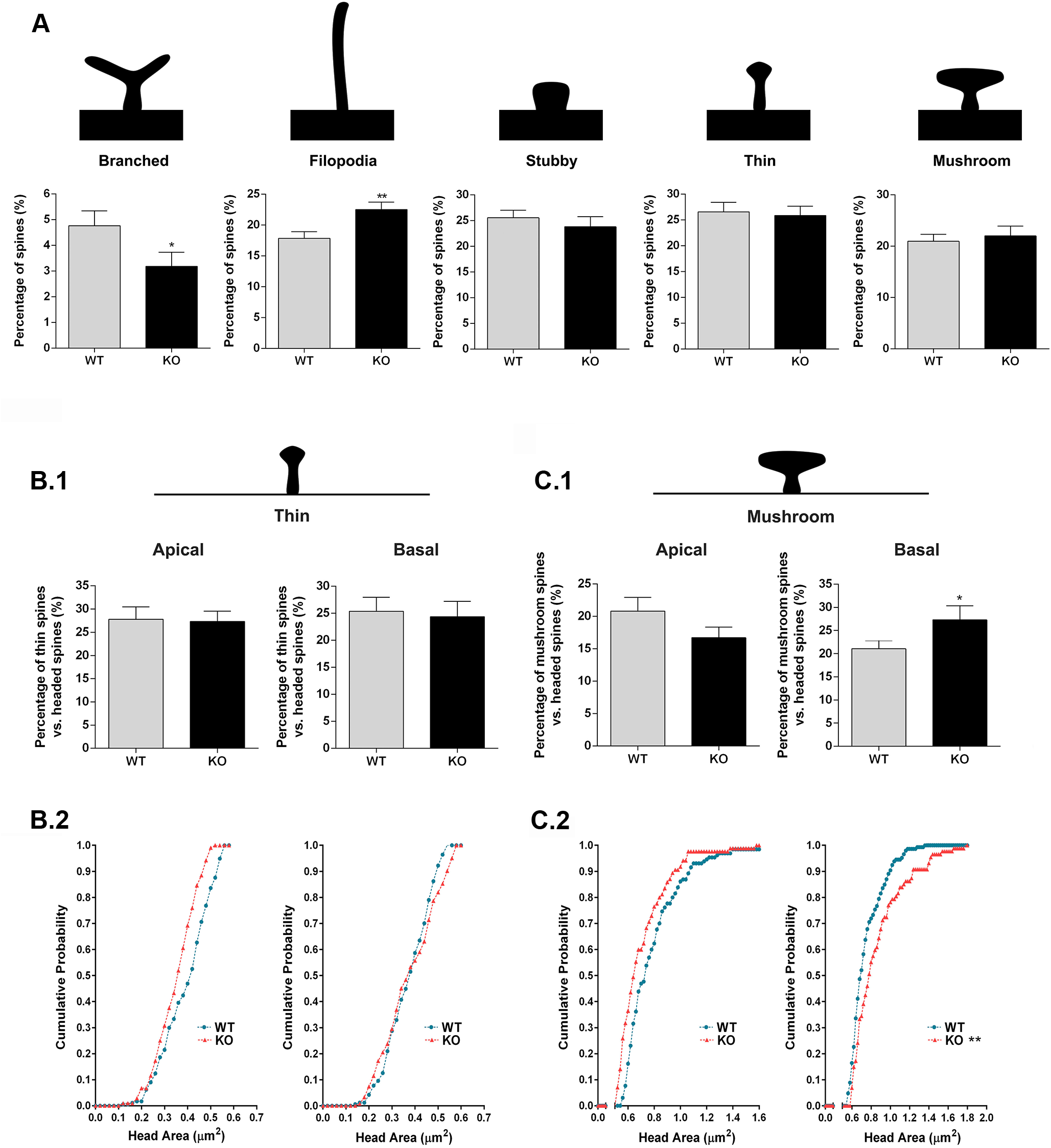
RTP801 modulates spine morphology of pyramidal neurons in the motor cortex of trained animals. Golgi-stained brains were processed and spine morphology of pyramidal neurons from M1 layer V were analyzed. Schematic representations show the spine morphologies considered in this study. **A.** RTP801 KO mice show reduced percentages of branched spines but increased percentage of filopodia spines in layer V neurons. Graphs show the percentage of each morphological type of dendritic spines *versus* total number of spines analyzed. Percentage of apical and basal thin (**B.1**) and mushroom (**C.1**) spines *versus* total number of spines with head. Spine density measures are represented as mean ± SEM of 50 dendrites per genotype (5 animals per genotype, 5 basal and 5 apical dendrites per animal). Apical and basal spines were analyzed separately. Data in **A**, **B**.1 and **C**.1 was analyzed with two-tailed Student’s t-test, * *P*< 0.05, ** *P*< 0.01 *versus* WT. Cumulative probability of apical and basal spine head area in thin (**B.2**) and mushroom (**C.2**). Distributions were compared with the Kolmogorov–Smirnov test. Apical and basal spines were analyzed separately. 5 animals/genotype were analyzed, 5 apical and 5 basal dendrites per animal. Spine head area from basal mushroom spines shows significant differences between genotypes, D=0.2747, *P*=0.0021.

We next asked whether this evidence in LV pyramidal neurons based on Golgi-Cox staining could be supported ultra-structurally by Transmission Electron Microscopy (TEM). Interestingly, we observed that KO mice synapses had bigger postsynaptic area (around 10%) (**Fig 6 A**) along with a wider PSD area, length and thickness (around 5%, each) (**Fig 6 B**). Interestingly KO mice exhibited a higher percentage of contacts containing mitochondria, mostly at the presynaptic compartment although the postsynaptic compartment showed a similar tendency (**Fig 6 C**). We did not find significant differences in the percentage of presynapses with more than one post-synapse, postsynapses with more than one presynapse or postsynapses with spine apparatus (**Fig S5 A-C**). Altogether, these results support the idea that although the KO mice have a decreased number of spines in the motor cortex LV, they displayed a more efficient synaptic structure, leading to an improvement of motor learning skills.

**Figure 6.**
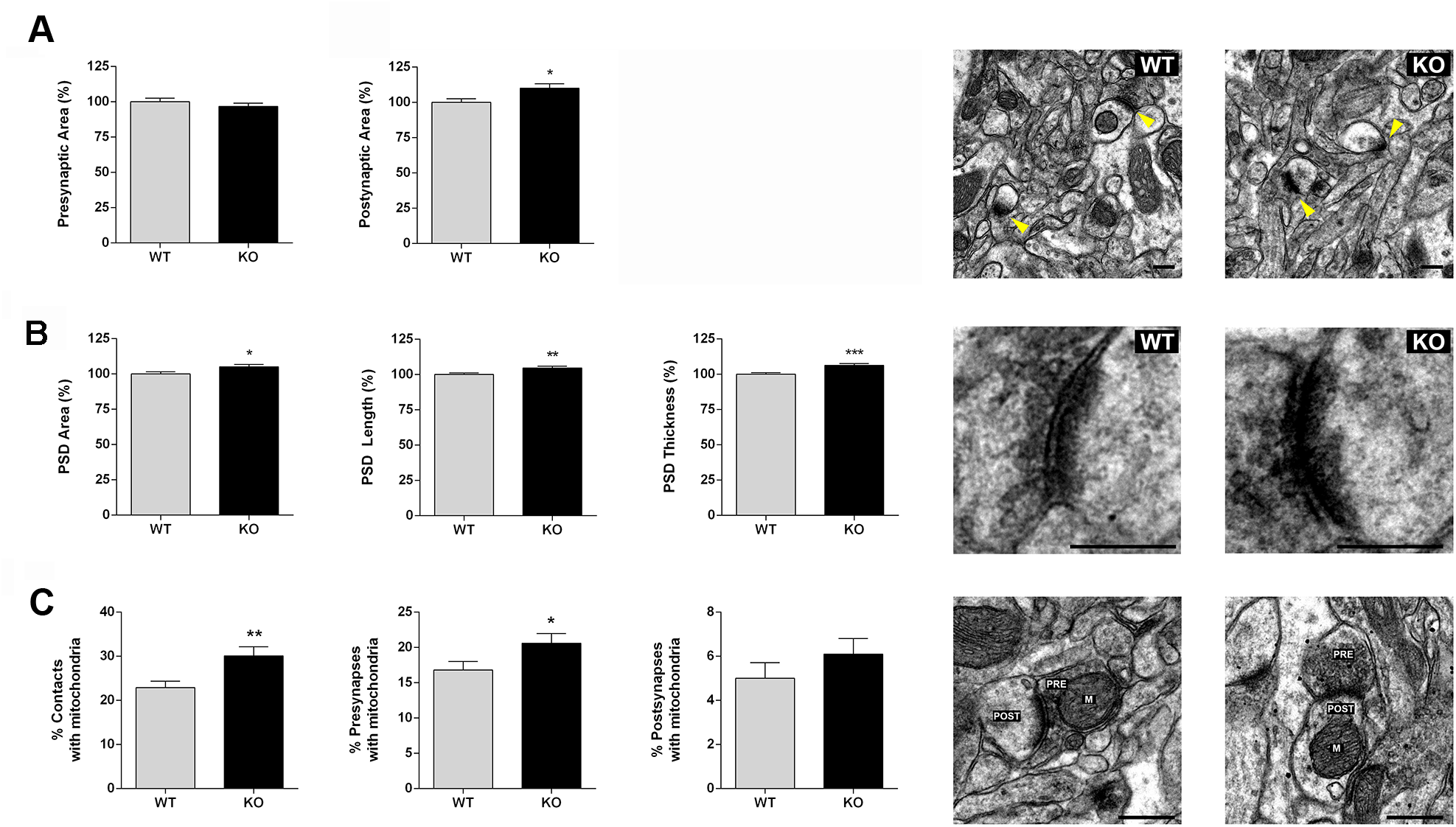
Synaptic contacts show structural differences between WT and RTP801 KO animals in the motor cortex after motor skill training. Motor cortex lower layers were analyzed by TEM**. A. Postsynaptic area is increased in RTP801 KO mice synaptic contacts. B. Postsynaptic density (PSD) area, length and thickness are increased in RTP801 KO mice synaptic contacts.** Histograms show mean ± SEM of the different measures analyzed relative to control mean. Images show a representative PSD for each genotype. **C. Increased presence of mitochondria in RTP801 KO synaptic contacts.** Graphs show the percentage of mitochondria found in synaptic contacts (either in the pre- or postsynaptic compartment) and in the presynapses or postsynapses separately. Data is represented as mean percentage ± SEM. Images show representative contacts where a mitochondrion (M) is present in either the presynapse (PRE) or postsynapse (POST). All histograms represent data from 45-50 images per animal, four animals per genotype. Data in a-c represent data relative to WT mean. Data was analyzed with Mann-Whitney test. * *P*< 0.05, ** *P*< 0.01, *** *P*< 0.001 *versus* WT control. For all electron micrographs, *Scale bar*, 250 nm.

### The lack of RTP801 elevates GluA1 AMPAR post-synaptically

In line with the reduction in spine density in neurons from motor cortex LV in the RTP801 KO mice (**Fig 4B**), biochemical analysis of KO motor cortex crude synaptic fractions confirmed a decrease in PSD-95 (**Fig 7 A**) but an specific enrichment of synaptic GluA1 (**Fig 7 B**), a crucial AMPAR subunit that has been described to be a key mediator in the acquisition of new motor skills (Kida *et al*, 2016; Roth *et al*, 2020). On the other hand, GluA2 AMPAR subunit, the prototypical auxiliary subunit of AMPARs stargazin or the N-methyl-D-aspartate receptor (NMDAR) subunit GluN2B did not change in KO mice in comparison to WT (**Fig 7 B-C**). Interestingly, we observed that levels of TrkB were also elevated in total homogenates in the RTP801 KO motor cortex (**Fig 7 D**), supporting the idea of a synaptic strengthening. By immunostaining WT and KO sections against PSD95 and GluA1 postsynaptic markers, we confirmed these initial biochemical observations specifically in M1 layer V. Indeed, the number of PSD-95 and GluA1 puncta diminished in the KO animals (**Fig 7E, H**) although the area and the intensity of the GluA1 dots were increased (**Fig 7 F-G**). Area and intensity of PSD-95 positive dots showed a non-significant increased tendency, as well (**Fig 7 F-G**). Altogether, these results suggest a novel synaptic role for RTP801 modulating synaptic strength and motor learning in the motor cortex (**Fig 8**).

**Figure 7.**
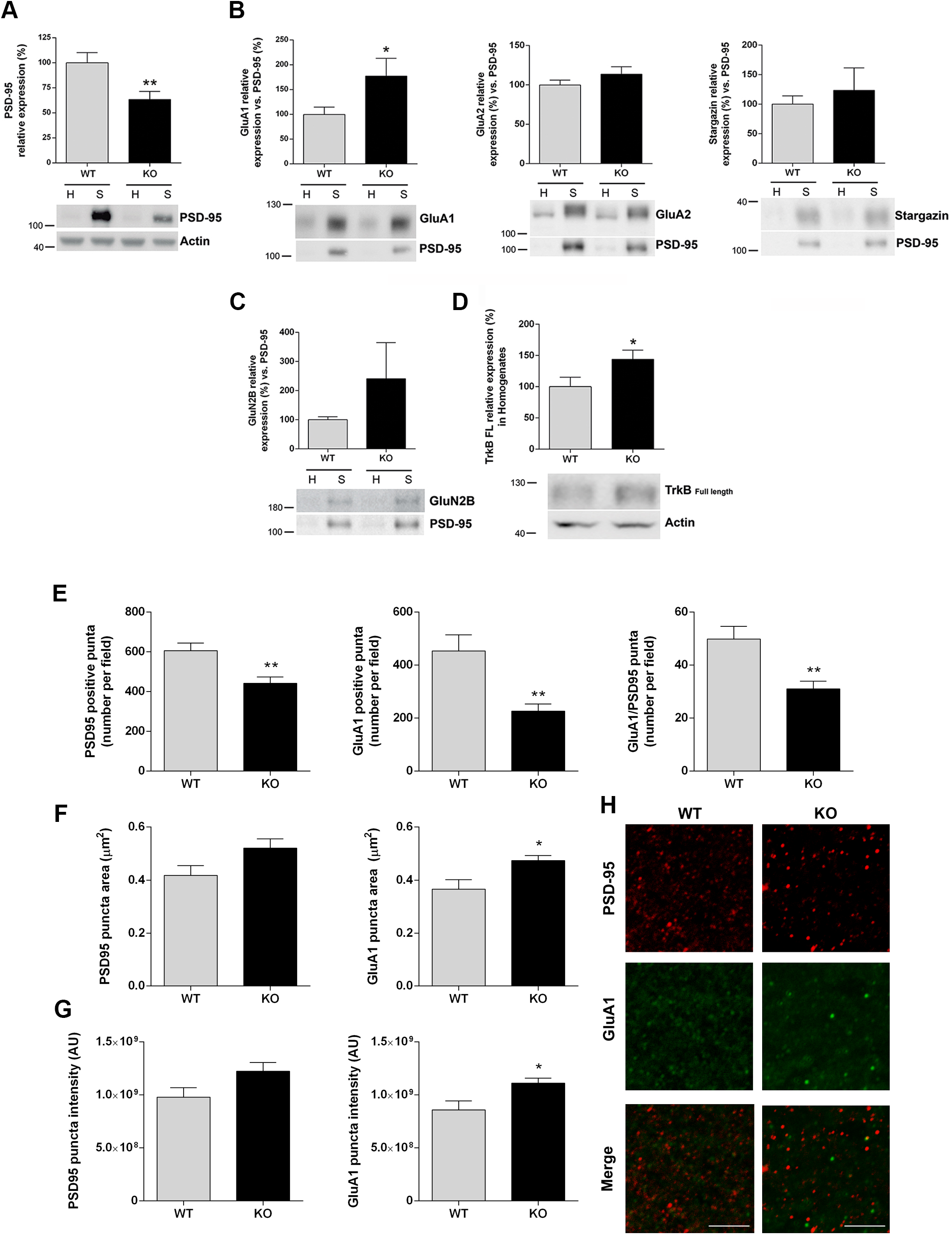
RTP801 modulates synaptic composition in the cortex from motor-trained mice. **A-C. Lack of RTP801 expression decreases PSD-95 levels but increases AMPAR subunit GluA1 levels in synaptosomes.** Levels of postsynaptic proteins were analyzed in 2-months-old WT and RTP801 KO animals. Crude synaptosomal fractions were obtained from cortical brain lysates and analyzed by western blotting. Representative images of PSD-95 (**A**), GluA1, GluA2, Stargazin (**B**), GluN2B (**C**) and actin are shown. **D. Lack of RTP801 expression increases TrkB receptor levels in motor cortex homogenates.** In the same WT and KO samples levels of full length TrkB was assessed by western blotting from homogenates. Representative images of TrkB and actin are shown. Densitometric measures (mean ± SEM) of total levels of PSD-95 and TrkB were relativized against actin and synaptic levels of the different proteins in the synaptic fraction were relativized against synaptic marker PSD-95. **E-H.** M1 LV excitatory postsynaptic characterization in 2-months old WT and RTP801 KO mice after performing behavioral motor tasks. **E.** Quantification of the number of PSD-95, GluA1 and PSD-95/GluA1 positive puncta per field. **F-G**. Quantification of PSD-95 and GluA1 puncta mean area (**F**) and intensity (**G**). Representative confocal images of a double immunofluorescence of PSD-95 and AMPAR subunit GluA1 in the motor cortex layer 5. *Scale bar*, 100 μm. All values appear as mean ± SEM and were analyzed with two-tailed Student’s t-test *versus* WT * *P*< 0.05 and **P<0.01.

**Figure 8.**
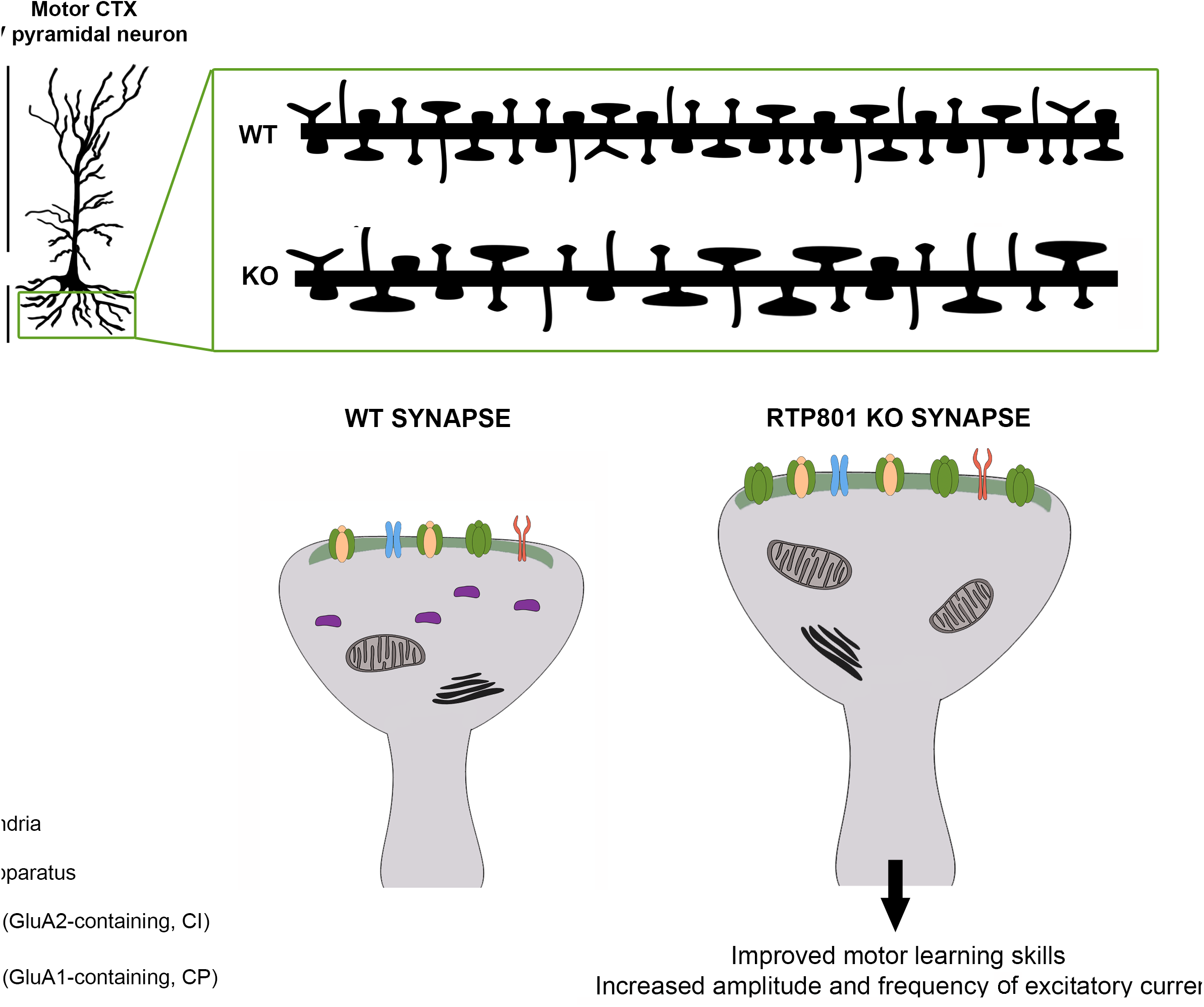
RTP801 KO mice show improved motor learning skills accompanied by functional and structural differences at a synaptic level. In comparison to WT mice, RTP801 KO mice show decreased spine density in M1 LV neurons together with an increase in the proportion of filopodia and mushroom-like dendritic spines. At a structural level, we found increased post-synaptic areas and PSD size and increased presence of mitochondria at the synapse in KO primary motor cortex LV together with increased levels of synaptic GluA1 AMPAR subunit.

## DISCUSSION

Here, we show a novel role for RTP801 in the modulation of synaptic plasticity in motor learning. The lack of RTP801 in mice resulted in decreased spine density and enhanced synaptic transmission in the primary motor cortex together with a better performance in the accelerating rotarod but altered gait and clasping. This improvement in motor learning skills was associated with alterations in dendritic spine structure. Cortical neurons in the motor cortex M1 layer V showed higher number of filopodia- and a mushroom-like morphology and TEM analyses revealed increased postsynaptic size in neurons from LV. In line with that, trained RTP801 KO mice showed higher levels of synaptic AMPAR subunit GluA1 and a general increase in TrkB levels.

Since the only evidences that RTP801 could modulate synaptic plasticity were found in pathological conditions, here we studied for the first time the putative role of RTP801 in a physiological context. In a context of depressive disorders, RTP801 KO mice were found resilient to stress-induced synaptic loss in the PFC (Ota *et al*, 2014; Kabir *et al*, 2017). Moreover, RTP801 downregulation alleviated stress-induced neurodegeneration in a mouse model of genetic PD (Zhang *et al*, 2018). More recently, our group described that synaptic RTP801 mediated motor-learning dysfunction in the R6/1 mouse model of HD (Martín-Flores *et al*, 2020). However, its potential physiological synaptic role has never been investigated in depth.

Hence, we initially confirmed that RTP801 was present in the synapses from a wide range of human and murine samples, as we previously described in HD murine models and HD human postmortem samples (Martín-Flores *et al*, 2020). Interestingly, RTP801 was highly enriched in human and rat crude synaptosomes but not that elevated in synaptic WT mice samples. In line with that, in cortical cultures RTP801 was localized mostly post-sinaptically. Interestingly, we found that spine density decreased in cortical cultures when RTP801 expression was transiently downregulated and that was translated with an increase in the amplitude and frequency of mEPSCs in KO cortical cultures. An opposite effect was found when ectopic RTP801 was expressed in hippocampal primary cultures.

Previous studies pointed out that RTP801 KO mice had normal brains and similar behavior to WT animals (Brafman *et al*, 2004; Ota *et al*, 2014). However, no thorough behavioral, biochemical and histological studies were performed in these animals. Macroscopically, we found that the KO mice brain weight less than WT brains, independently of the total body size, and it was likely due to a decrease in the cell density of M1 LV. Noteworthy, this difference can be explained by the developmental role of RTP801, which regulates both neurogenesis by regulating neuroprogenitors’ proliferation rate and neuronal migration/differentiation in the cortex (Malagelada *et al*, 2011).

*In vivo*, the lack of RTP801 reduced spine density in the M1 layer V in the KO mice vs. WT. We observed a similar result when we transiently downregulated RTP801 in cultured cortical neurons (**Fig 1C**). Interestingly, KO animals showed higher synaptic performance in KO motor cortex (LV) slices *versus* WT. These results therefore suggest that the lack of RTP801 decreases spine density but enhances synaptic function.

To investigate the role of RTP801 in synaptic plasticity *in vivo* we performed several motor behavioral tests and checked circuitries that control movement and motor learning. RTP801 KO mice showed gait impairment but no alterations in general locomotor activity. It is noteworthy that gait abnormalities are more likely to be explained by cerebellar dysfunction and more studies will be needed in the future. Despite gait alterations, the lack of RTP801 improved mouse motor learning skills. These results are in line with the work of Zhang et al (2018) (Zhang *et al*, 2018), where the knockdown of RTP801 in the substantia nigra partially rescued motor function in a mouse model of PD subjected to chronic-restraint stress. In addition, we recently described that striatal RTP801 knock down in the R6/1 mouse model of HD prevented from motor-learning deficits (Martín-Flores *et al*, 2020).

The most characterized circuitry involved in motor learning is the corticostriatal pathway. Pyramidal neurons from the M1 along with striatal MSNs predominantly undergo synaptic dynamics under motor learning (Tjia *et al*, 2017; Costa *et al*, 2004). Indeed, spine density in the M1 LV neurons from the RTP801 KO mice, specifically in their basal dendrites, was decreased. We did not observe any differences in spine density in the apical dendrites of the same neurons or in the striatal MSNs from the KO mice. Related to the cortex, Ota and colleagues (Ota *et al*, 2014) did not find spine density differences in the prefrontal cortex (PFC) between WT and RTP801 KO mice in basal conditions. This fact, together with the absence of differences in the striatum in our work, may point towards a region-specific role of RTP801 in the normal (or physiological, non-stressed) mouse brain. Hence, RTP801 could be contributing to motor learning at the basal dendrites of LV pyramidal neurons.

This speculation was also supported by the observations of the synaptic morphology in the motor cortex LV pyramidal neurons, where we observed a significant increase of filopodia along with a decrease in branched spines in the KO animals. Although the physiological meaning of branched spines is still in debate, filopodia have been proposed to be precursors of spines, to develop an explorative role to increase the probability to form a synapse (Ziv & Smith, 1996; Zuo *et al*, 2005). Thus, an increase in this type of spines could explain the better performance of the KO mice in the accelerating rotarod. However, filopodia-related plasticity must have a fine-tuned regulation, since a high remodeling rate might be troublesome (reviewed in (Ozcan, 2017)). Indeed, among headed spines, we detected an increase in the percentage and head area of mushroom-like spines from basal dendrites between WT and KO animals. This fact correlates well with the change of spine morphology and the function of the spines, and in the end, with an increase in synaptic strength of the area (Arellano *et al*, 2007; Yuste *et al*, 2000).

We confirmed a more complex postsynaptic compartment by TEM. The lack of RTP801 led to an increase in postsynaptic area in the synapses of the region of study, although no differences were detected in the presynaptic compartment. Strikingly, greater postsynaptic density size was detected in RTP801 KO animals in the same area. Interestingly, a positive correlation between the amount of PSD and spine size (Arellano *et al*, 2007) and the former with synaptic strength (Béïque & Andrade, 2003; Meyer *et al*, 2014) has been described. Moreover, KO synaptic contacts present more mitochondria, whose presence at the synapse has been related with a role in controlling plasticity processes (Todorova & Blokland, 2017; Lee *et al*, 2018)). Our ultrastructural analyses, therefore, seem to indicate that, although the lack of RTP801 causes a decrease in spine density, the remaining spines are able to compensate this reduction at a structural level.

Interestingly, we observed a differential synaptic composition in the remaining spines in the M1 LV from RTP801 KO mice versus WT animals. We observed decreased levels of synaptic PSD-95 in crude synaptosomes that go in line with the decreased number of PSD-95 positive puncta in M1 LV observed by immunohistochemistry, along with a specific elevation of GluA1 AMPAR subunit at the synapses in M1 LV of KO mice. Calcium impermeable AMPARs (GluA2-containing; CI-AMPARs) are the most prevalent type of AMPAR in neurons (Lu *et al*, 2009) where they are responsible for postsynaptic currents and the depolarization of the postsynaptic neuron. In contrast, GluA1 subunit confers calcium permeability to the receptor. Calcium permeable AMPAR (CP-AMPARs) are mostly engaged to synaptic regulation and intracellular signaling (reviewed in (Man, 2011). Therefore, the improved performance observed in the KO mice could be explained at least in part with this change in the AMPA receptors subunit composition. This could favor the presence of CP-AMPARs with high calcium permeability and then, in consequence, signaling activation and synaptic regulation. Interestingly, previous studies have demonstrated that motor learning induces an increase in GluA1 levels in dendritic spines in the motor cortex. This increment in GluA1 subunits are key modulators of synaptic plasticity induced by motor skill learning (Roth *et al*, 2020). The mechanism by which RTP801 could mediate this specific AMPAR subunits composition at the synapses to modulate motor learning has to be explored yet. Ectopic RTP801 overexpression showed the opposite result, since it reduced GluA1 puncta intensity in cultured hippocampal neurons. Remarkably, RTP801 silencing in R6/1 mice induced an increase of total levels of GluA1 and TrkB neurotrophin receptor. Indeed, in trained RTP801 KO mice we could also observe an increase in total levels of TrkB receptor.

This result is in line with other works describing that synaptic activity modulates both BDNF levels and TrkB receptors amount and localization (Guo *et al*, 2014; Lauterborn *et al*, 2000).

In summary, our work indicates a novel synaptic function for RTP801 in motor learning by modulating synaptic structure, composition and plasticity. This finding is important since motor learning impairment is a key feature of neurodegenerative diseases such as PD and HD. Altogether, our results point towards RTP801 downregulation as a promising therapeutic strategy to ameliorate motor learning dysfunction in these diseases.

## MATERIALS AND METHODS

### Animals

Transgenic RTP801 knock out mouse strain was generated by Lexicon Inc. as described in (Brafman *et al*, 2004). RTP801 knockout mice were obtained by homozygous pairing. Thus, wild type mice were bred from the RTP801 KO founder strains to obtain a C57Bl6/129sv background. RTP801 knock out and wild type mice were housed under controlled conditions (22°C, 40-60% humidity in a 12-hour light/dark cycle) with water and food available *ad libitum*. All the animals analyzed in this study were 2 months-old adult mice.

For further biochemical analyses, Golgi staining and TEM, mice were euthanized by cervical dislocation and tissue was dissected out. For immunohistochemistry, animals were processed as described elsewhere (Creus-Muncunill *et al*, 2018). Briefly, animals were anesthetized with 60mg/kg dolethal and intracardially perfused with 4% PFA. Coronal 25μm-thick brain sections were obtained with a cryostat.

### Rat primary cultures

Rat cortical and hippocampal primary cultures were obtained from embryonic day 18 Sprague-Dawley rats as previously described (Canal *et al*, 2016). Cells were either transduced with lentiviral particles carrying a control shRNA or a specific shRNA against RTP801 or transfected with lipofectamine 2000 (Thermo Fisher Scientific) with pCMS vectors expressing eGFP (donated by Dr. Lloyd Greene, Columbia University) or eGFP-fused RTP801 protein (Romaní-Aumedes *et al*, 2014). The sequences to downregulate or overexpress RTP801 were previously described in (Malagelada *et al*, 2006).

### Mouse primary cultures

Mouse primary cortical cultures were obtained from embryonic day 15 mice. Coverslips were coated for 1h with 0.1 mg/ml poly-D-lysine (Merck) and then 3.5h with 0.018 mg/ml laminin (Thermo Fisher Scientific). Briefly, cortices were dissected out and chemically digested with 41.66μM Trypsin for 10 minutes. Following mechanical digestion, cells were plated on coverslips at a density of 25.000 cells/cm^2^ and maintained in Neurobasal-A medium supplemented with B27, GlutaMAX (all from Gibco), 33.3 mM Glucose and 1% penicillin-streptomycin (Sigma) in a 5% CO_2_ atmosphere and 37°C.

### Crude synaptosomal fractionation

Tissue (rat, mice or *postmortem* human brains) or cultured cells were homogenized in Krebs-Ringer buffer (125mM NaCl, 1.2mM KCl, 22mM NaHCO3, 1mM NaH2PO4, 1.2mM MgSO4, 1.2mM CaCl2, 10 mM Glucose, 0.32 M Sucrose; pH 7.4). For samples in Figure 7, mice were sacrificed one week after behavioral testing. Initial lysate was first centrifuged at 1.000g for 10 minutes. Supernatant (homogenate) was centrifuged for 20 minutes at 16.000g to obtain the cytosolic fraction (supernatant) and the crude synaptosomal fraction (pellet), that was resuspended in Krebs-Ringer buffer.

### Western blotting

Samples were resolved in NuPAGE™Novex™ polyacrylamide gels and proteins were transferred to nitrocellulose membranes with the iBlot system (all from Thermo Fisher Scientific). Indicated primary antibodies were incubated overnight at 4°C diluted in Tris-buffered saline containing 0.1% Tween-20 and 5% BSA. Secondary antibodies (Thermo Fisher Scientific) were diluted in TBS-Tween with non-fat dry 5% milk (Bio-rad) for 1 hour. Proteins were detected with Supersignal™ West Pico Plus chemiluminiscent substrate (Thermo Fisher Scientific) and images were acquired with ChemiDoc™ (Bio-Rad). The following antibodies were used: RTP801 (1:500, Proteintech), HRP-conjugated anti-beta actin (1:100.000; Sigma), PSD-95 (1:1000; Thermo Fisher Scientific), SV2a and GFP (1:1000; Santa Cruz Biotechnology), GluA1, GluA2, Stargazin (1:1000; Merck Millipore), GluN2B (1:1000; Cell Signaling Technology) and TrkB (1:1000; BD Biosciences).

### Immunofluorescence

Cells were fixed in 4% PFA and permeabilized with 0.25% Triton-X. Blocking and antibody incubation was performed with Superblock (Thermo Fisher Scientific). Primary antibodies were incubated over night at 4°C and secondary antibodies for 2h at room temperature. For mouse brain tissue immunofluorescence, sections were washed with PBS and incubated for 30 min in NH_4_Cl. Next, sections were blocked with 0.3% Triton-100 10% NGS in PBS for 2h prior incubation with the primary antibodies diluted in blocking solution overnight at 4°C. Later, sections were washed and incubated for 2h with the secondary antibodies. Slices were then washed with PBS. Both cells and tissue samples were mounted with Prolong Gold antifade mountant (Thermo Fisher Scientific). The following antibodies were used: GFP (1:500), SV2a (1:100) (both from Santa Cruz Biotechnology), PSD-95 (1:50; Thermo Fisher Scientific), GluA1 (1:250-1:500; Merck Millipore) and RTP801 (1:100; Proteintech). AlexaFluor-488 or −555 secondary antibodies (1:500) and Hoechst33342 (1:5000) were from Thermo Fisher Scientific. Images were obtained with a Leica LCS SL or a Zeiss LSM880 confocal microscopes with a 1024×1024 pixel resolution and a 63x magnification and were analyzed with ImageJ. For in vitro experiments in cortical neurons, at least 25 dendrites per group from three independent experiments were analyzed. For in vitro experiments in hippocampal neurons, at least 12 neurons per group were analyzed from three independent experiments. For double-labeled GluA1-PSD-95-positive clusters in brain slices, images were acquired with 4x digital zoom (33.74×33.74 μm). For each mouse three representative images from two different coronal sections were analyzed. Colocalization was considered when there was at least one common pixel between GluA1 and PSD-95 detected puncta.

### Nissl staining

Slices were stained for 45 min with 0.2 mg/ml Cresyl violet (Sigma) in a 0.1 M acetic acid 0.1 M sodium acetate solution. Next, slices were washed in distilled water and then dehydrated with ethanol (70, 95, 100%, 5 minutes each), washed with xylol and mounted with DPX media. Images were obtained with a 10x magnification with a Zeiss Axiolab.

### Behavioral assessment

#### Footprint test

Mice’s fore and hindlimbs were painted in blue and red, respectively, with non-toxic ink. Animal’s gait was then recorded letting them walk through a tunnel on white paper (10 cm wide, 40 cm long). The test was performed three times on the same day. In each trial three consecutive steps were measured for each parameter (stride, sway, stance, overlap).

Open field test: mice were placed in a 40×40×40 cm arena. The center area was considered as the central squared 20×20 cm space. Light intensity was 24 lux though-out the periphery and 29 lux in the center. Mice’s movement was tracked and recorded for 10 minutes using SMART 3.0 Software (Panlab). Other parameters related to anxiety-like behaviors, like number of groomings, rearings and defecations were also monitored.

#### Accelerating rotarod

one day after the Open field test mice were subjected to the Accelerating rotarod test. Mice were placed on a 3 cm rod with an increasing speed from 4 to 40 rpm over 5 minutes. Latency to fall was recorded as the time mice spent in the rod before falling. Accelerating rotarod test was performed for 4 days, 4 trials per day. Trials in the same day were separated by 1 hour.

#### Clasping behavior

Hindlimb clasping was measured by picking up mice at the base of the tail. In order to classify this phenotype we used the scale described in (Guyenet *et al*, 2010) with minor modifications: 0 means no hind paw retraction, 1, one hindlimb retracted, 2, both hindlimbs partially retracted, and 3 when the 2 hindlimbs were totally retracted.

### Golgi Staining and spine density and morphology analyses

Golgi-Cox impregnation was performed with fresh brain hemispheres from, mice sacrificed one week after behavioral testing with FD Rapid GolgiStain™kit (FD Neurotechnologies) following manufacturer’s instructions. 100 μm slices were obtained with a Leica vibratome and mounted on gelatin-coated slides before final staining.

For spine density analyses only pyramidal neurons from layer V in the primary motor cortex or medium spiny neurons (MSNs) from the dorsolateral striatum were taken into account. Spine density was quantified in dendritic segments of at least 10 μm and 30 different secondary/tertiary dendrites per animal were analyzed. Analyzed dendrites were 50% apical, 50% basal.

Spine morphology analyses were performed in motor cortex layer V pyramidal neurons. Spines in 5 apical and 5 basal secondary/tertiary dendrites were analyzed for each animal (6 WT and 4 KO), in segments of at least 10 μm long. A total of 100-125 apical and 100-125 basal spines were analyzed per animal. Branched, filopodia and stubby spines were visually categorized. For headed spines, head area was measured in all headed spines and thin/mushroom classification was performed depending on the mean head area for each genotype (spines with head area greater than the mean were considered as mushroom spines and smaller ones were categorized as thin spines). In spine density and morphology analyses, animal genotype was blind for the experimenter.

### Transmission electron microscopy

2 months old RTP801 knock out (n=4) and wild type mice (n=4) were sacrificed one week after behavioral testing and motor cortex was dissected from coronal sections. From these sections, the lower half of the motor cortex, including Layer V and VI, was isolated and fixed overnight in 2% glutaraldehyde 2% paraformaldehyde in 0.12 M phosphate buffer. After fixation, tissue was processed and analyzed as previously described in (Bosch *et al*, 2016). Electron micrographs were randomly taken at 25.000x with a TEM JEOL J1010 (tungsten filament), with a CCD Orius (Gatan) and software Digital Micrograph (Gatan). Spine density, pre/postsynaptic area and postsynaptic density area, length and thickness were determined (n=45-50 images for each animal) with ImageJ software. In all TEM analyses, animal genotype was blind for the experimenter.

### Electrophysiology

#### Rat neuronal cultures

miniature excitatory postsynaptic currents (mEPSCs) were measured in rat primary hippocampal neurons plated on glass coverslips as previously described (Gilbert *et al*, 2016).

#### Mouse cortical cultures

Electrophysiological recordings of cultured cortical pyramidal neurons –chosen in basis of their characteristic pyramidal morphology– were performed at 14 DIV. Whole-cell patch-clamp currents were recorded at room temperature (25–26 °C) in extracellular solution containing (in mM): 130 NaCl, 3.5 KCl, 10 HEPES, 15 glucose and 2 CaCl_2_ (pH 7.4; osmolarity 305 mOsm/Kg with sorbitol). AMPAR-mediated miniature excitatory postsynaptic currents (mEPSCs) were isolated adding to the extracellular solution 1μM tetrodotoxin to block evoked synaptic transmission, 100μM picrotoxin to block GABA_A_ receptors and 50μM APV to block NMDA receptors. Recording electrodes were fabricated from borosilicate glass with a final resistance of 4–5 MΩ and filled with an internal solution containing (in mM): 120 K-Gluconate, 16 KCl, 8 NaCl, 10 HEPES, 0.2 ethylene glycol tetraacetic acid (EGTA), 2 MgATP, 0.3 Na_2_GTP (pH 7.2; osmolarity 291 with sorbitol). Recordings were acquired at a sampling rate of 5KHz and were filtered at 2Hz. Miniature events were detected and analyzed with the WaveMetrics Igor Pro open-source software package Neuromatic (Rothman & Silver, 2018). Frequency was determined by dividing the number of detected events by the recorded time (in seconds).

### Electrophysiological field recordings

Two-month old (female and male) mouse brain sagittal sections were obtained on a vibratome (Microm HM 650 V, Thermo Scientific, Waltham, MA, USA) at 350 μm thickness in oxygenated (95% O_2_, 5% CO_2_) ice-cold aCSF and then transferred to a oxygenated 32°C recovery solution for 15 min as previously described (Choi *et al*, 2019). Then, slices were transferred to oxygenated aCSF at room temperature and left for at least 1 h before electrophysiological field recording. Following recovery, mouse 350 μm thick brain slices were placed in a multi electrode array (MEA) recording dish and fully submerged in oxygenated aCSF at 37 °C. Electrophysiological data were recorded with a MEA set-up from Multi Channel Systems MCS GmbH (Reutlingen, Germany) composed of a 60 channels USB-MEA60-inv system. Experiments were carried out with 60MEA200/30iR-ITO MEA dishes consisting of 60 planar electrodes (30 μm diameter) arranged in an 8×8 array and placed in the motor cortex slice surface. Raw traces were recorded for 5 min from 58 electrodes simultaneously, sampled at 5 kHz. Raw data were high-pass filtered with a 200-Hz Butterworth 2^nd^ order filter, the noise level calculated by the standard deviation of the recorded signal on each electrode and spikes were identified as currents with a negative amplitude larger than −30 mV and slope values between 0.2 and 1. To quantify burst activity in spike-trains we applied the MaxInterval Method (Legendy & Salcman, 1985) with the following parameter values: maximum beginning ISI, 200 ms; maximum end ISI, 200 ms; minimum interburst interval, 20 ms, minimum burst duration 20 ms; minimum number of spikes in a burst, 5. Software for recording and signal processing was MC Rack from Multi Channel Systems. Using a digital camera during recording assessed the position of the brain slices on the electrode field to analyze information from electrodes specifically positioned on cortical layer V (**Fig 2 H**).

### Experimental design and statistical analyses

Graphs show results reported as mean+SEM. Data was assessed for normality using D’Agostino-Pearson, Shapiro-Wilk or Kolmogorov-Smirnov. Statistical analyses were performed using unpaired, two-tailed Student’s T-test for normally distributed data, Mann-Whitney test for non-parametric data and Two-way ANOVA followed by Bonferroni’s *post-hoc* tests to compare multiple groups, as appropriate and indicated in the figure legends. Values of *P*< 0.05 were considered as statistically significant.

### Ethical Approval and Consent to participate

All procedures were performed in compliance with the NIH Guide for the Care and Use of Laboratory Animals and approved by the local animal care committee of Universitat de Barcelona following European (2010/63/UE) and Spanish (RD53/2013) regulations for the care and use of laboratory animals.

Human samples were obtained following the guidelines and approval of the local ethics committee (Hospital Clínic of Barcelona’s Clinical Research Ethics Committee).

## ACKNOWLEDGEMENTS

The authors thank Dr. Sílvia Ginés, Dr. Verónica Brito and Dr. Albert Giralt for helpful discussion. We also thank Dr. Albert Martínez from the Faculty of Biology from our same University, for his TEM assessment and guidance. We thank the Neurological Tissue Bank of the Biobanc-Hospital Clínic-IDIBAPS (Barcelona, Spain) and Dr. Ellen Gelpi for providing human tissue samples. We thank Maria Calvo from the Advanced Microscopy Unit, Scientific and Technological Centers, University of Barcelona, for their support and advice in confocal techniques.

## AUTHOR CONTRIBUTION

L.P-S., N.M-F., M.M., J.S., A.LL., J.R-A., J.S., G.C., M.C., D.S., X.G., J.A and C.M. have contributed in the conception and design of the study, acquisition and analysis of data and in drafting the manuscript and figures. G.C., E.G-G., N.S-F., S.F-G., J.P.G., M.J.R., H-Y.M., E.F. and D.W., have contributed in acquisition and analysis of data and in drafting the manuscript and figures.

## ABBREVIATIONS

(AD): Alzheimer’s disease
(HD): Huntington’s disease
(KO): knock out
(mTOR): mechanistic target of rapamycin
(LV): primary motor cortex (M1), motor cortex layer V
(mhtt): mutant huntingtin
(MEAs): microelectrode arrays
(mEPSC): miniature excitatory postsynaptic currents
(PD): Parkinson’s disease
(iPSC): induced pluripotent stem cells
(PSD): postsynaptic density
(TEM): transmission electron microscopy
(TSC1/2): tuberous sclerosis complex
(SNpc): substantia nigra pars compacta
(WT): wild type

## SUPPLEMENTARY FIGURE LEGENDS

**Supplementary figure 1. RTP801 is present in the postsynaptic compartment and its overexpression alters synaptic transmission and the levels of synaptic proteins in primary cultured hippocampal neurons. A. RTP801 protein colocalizes with postsynaptic marker PSD-95.** Cells were fixed at DIV 21 and endogenous RTP801 (green) and PSD-95 (red) were analyzed by immunofluorescence. White arrows designate PSD-95 and RTP801 colocalizations. *Scale bar*, 5 μm. **B. RTP801 seldom colocalizes with presynaptic protein SV2A.** Cells were fixed at DIV 17 and RTP801 (green) and SV2A (red) proteins were analyzed by immunofluorescence. White arrows point SV2A positive presynaptic *puncta*, which does not colocalize with RTP801. *Scale bar*, 5 μm. **C, D. Ectopic RTP801 reduces the intensity of PSD-95 and GluA1 at the synaptic contacts.** Neurons were transfected with a vector expressing RTP801 tagged with eGFP (eGFP-RTP801) or with an eGFP control vector (eGFP). Two days following transfection, at DIV 15, neurons were fixed and subjected to immunofluorescent staining against eGFP (green) and PSD-95 (**C**) or GluA1 (**D**) (both in red). Images show a representative staining for each protein. Graphs show immunofluorescent integrated intensity and synaptic accumulation density, represented as mean ± SEM from the analysis of at least 12 neurons per group. Data in c was analyzed with two-tailed Student’s t-test. Data in d was analyzed with Mann-Whitney test. * *P*< 0.05, ** *P*< 0.01 *vs*. eGFP. *Scale bar*, 10 μm. **F. Ectopic RTP801 attenuates postsynaptic excitatory transmission.** Primary cultures of hippocampal neurons from rat E.18 embryos were transfected at DIV 13 with a control vector (eGFP) or a vector expressing RTP801 protein fused with GFP (eGFP-RTP801). Two days later mEPSC were recorded through patch-clamp. Quantitative analysis of mEPSCs amplitude and frequency is represented in the graphics as mean ± SEM from the recordings performed in at least 8 transfected neurons per condition. Data was analyzed with Student’s t-test, * *P*< 0.01 *vs*. eGFP.

**Supplementary figure 2. Multielectrode array analysis of M1 LV spontaneous activity on mouse slices**. **A**. We found no sex effects in the spike rate differences between WT and RTP801 KO animals (Two-way ANOVA; F_(1,11)_ = 5.137; *P* = 0.045; F_(1,11)_ = 0.374; for the genotype effect; *P* = 0.553 for the sex effect, F_(1,11)_ = 1.627, *P* = 0.228 for the sex/genotype interaction). When we analyzed the spike-train patterns in WT and KO motor cortex slices we found no differences in mean burst duration (**B**), inter-spike interval duration (ISI) inside the bursts (**C**), and spike frequency in the burst (**D**).

**Supplementary figure 3. A. RTP801 KO mice show hindlimb clasping.** Example of clasping reflex; 2-months-old RTP801 KO mice display their hindlimbs towards their abdomen when suspended by the tail. Graphical representation of clasping phenotype scored: 0 (no clasping), 1 (one hindlimb is retracted), 2 (both hindlimbs are partially retracted) and 3 (both hindlimbs totally retracted towards the abdomen). Measures are represented as mean ± SEM and analyzed with two-way ANOVA (genotype effect, *** *P*< 0.001) followed by Bonferroni’s multiple comparisons test for *post hoc* analyses, *** *P*< 0.001. **B. Gait abnormalities are found in both male and female RTP801 KO mice.** Graphics show hindlimb lengths for stride, sway, stance and limbs overlap. Data is represented as mean ± SEM and was analyzed with two-way ANOVA followed by Bonferroni’s multiple comparisons test for *post hoc* analyses. * *P*< 0.05, ** *P*< 0.01, *** *P*< 0.001 *versus* same-gender WT group, N = 8 WT males, 5 WT females, 4 KO males and 8 KO females. **C. Anxiety-like behavior analyses during open field test.** The assessment was performed for 10 minutes. Graphs show measures of self-grooming behavior, number of wall and vertical rearings and number of fecal pellets. Data is represented as mean ± SEM and was analyzed with two-way ANOVA followed by Bonferroni’s multiple comparisons test for *post hoc* analyses. *** *P*< 0.001 *versus* same-gender WT group .N = 8 WT males, 5 WT females, 4 KO males and 8 KO females. **D. Both male and female RTP801 KO mice display the same tendency to perform better in the Accelerating Rotarod.** Time spent in the accelerating rotarod for three days, four trials per day, by RTP801 KO female and male mice. Data is represented as mean ± SEM and was analyzed with two-way ANOVA. N = 14 WT males + 17 WT females and 12 KO males + 18 KO females.

**Supplementary figure 4. Percentage of different types of spine morphology measured in apical and basal dendrites.** Graphs show the percentage of each morphological type of dendritic spines versus total number of spines. Measures are represented as mean ± SEM 50 dendrites per genotype (5 animals per genotype, 5 basal and 5 apical dendrites per animal). Data was analyzed with two-tailed Student’s t-test, * P<0.05.

**Supplementary figure 5. A-B.** There are no differences in the number of presynapses and postsynapses with more than one contact between WT and KO animals. Electron micrographs illustrate a presynapse (**A**) with two postsynaptic contacts and a postsynapse (**B**) with two presynaptic inputs. **C.** There are no differences in the percentage of postsynaptic compartments with spine apparatus (S. App) between genotypes. Image on the right show a perforated contact with a spine apparatus in the postsynapse. Data is represented as mean percentage ± SEM. All histograms represent data from 45-50 images per animal, four animals per genotype. Data was analyzed with Student’s t-test.

