## Supplementary figures and images for "RTP801 REGULATES MOTOR CORTEX SYNAPTIC TRANSMISSION AND LEARNING"

### S1

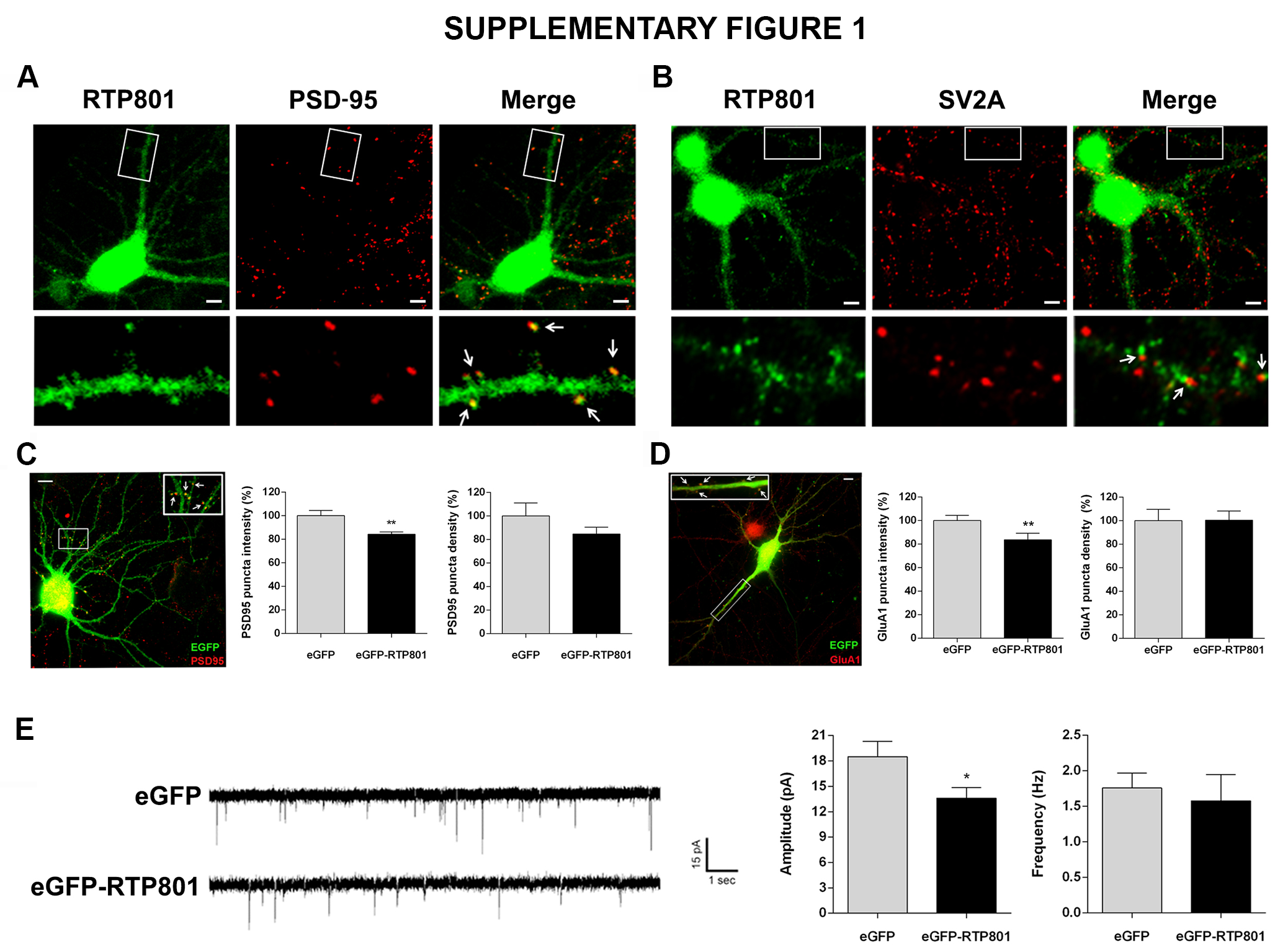

### S2

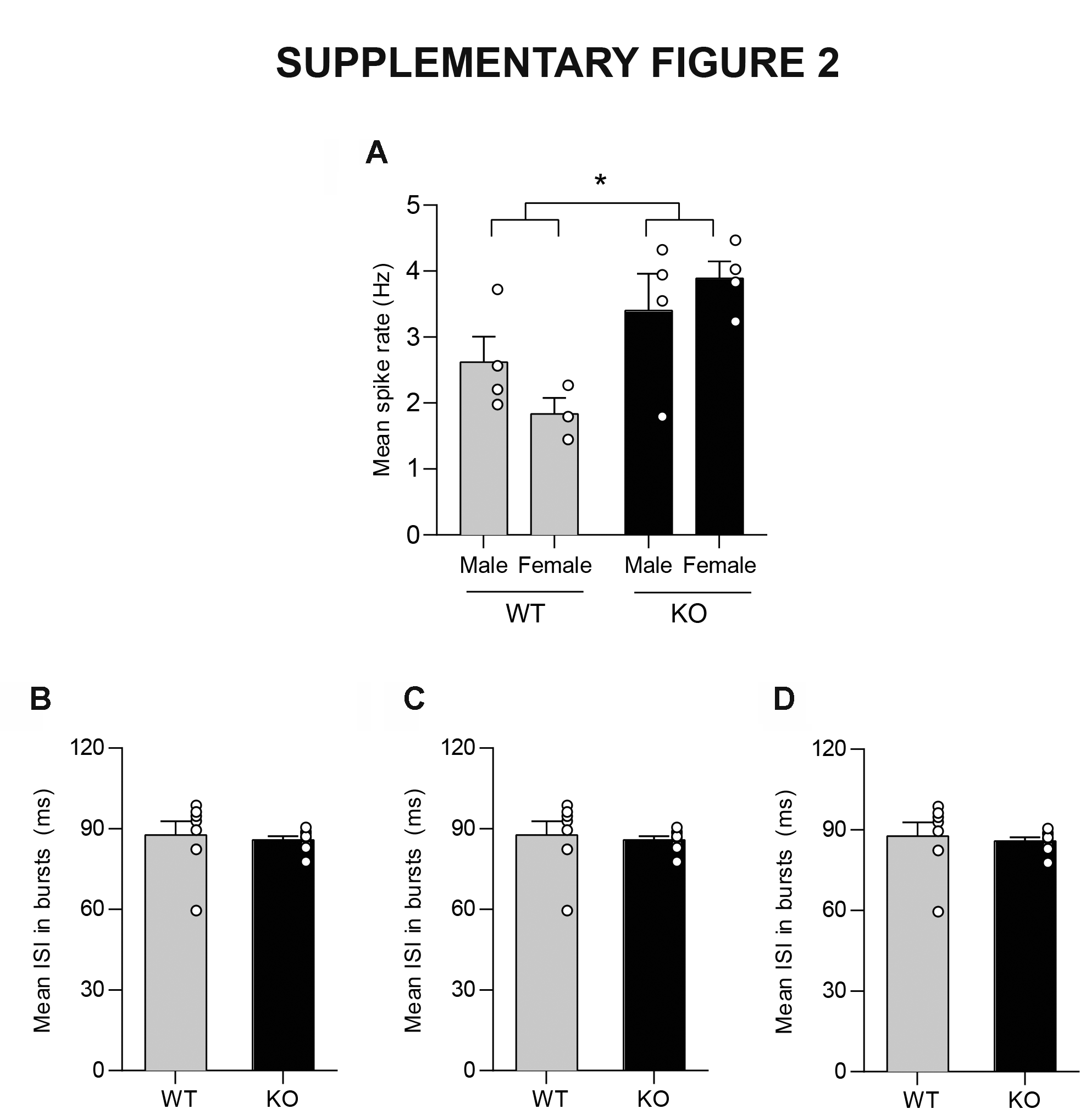

### S3

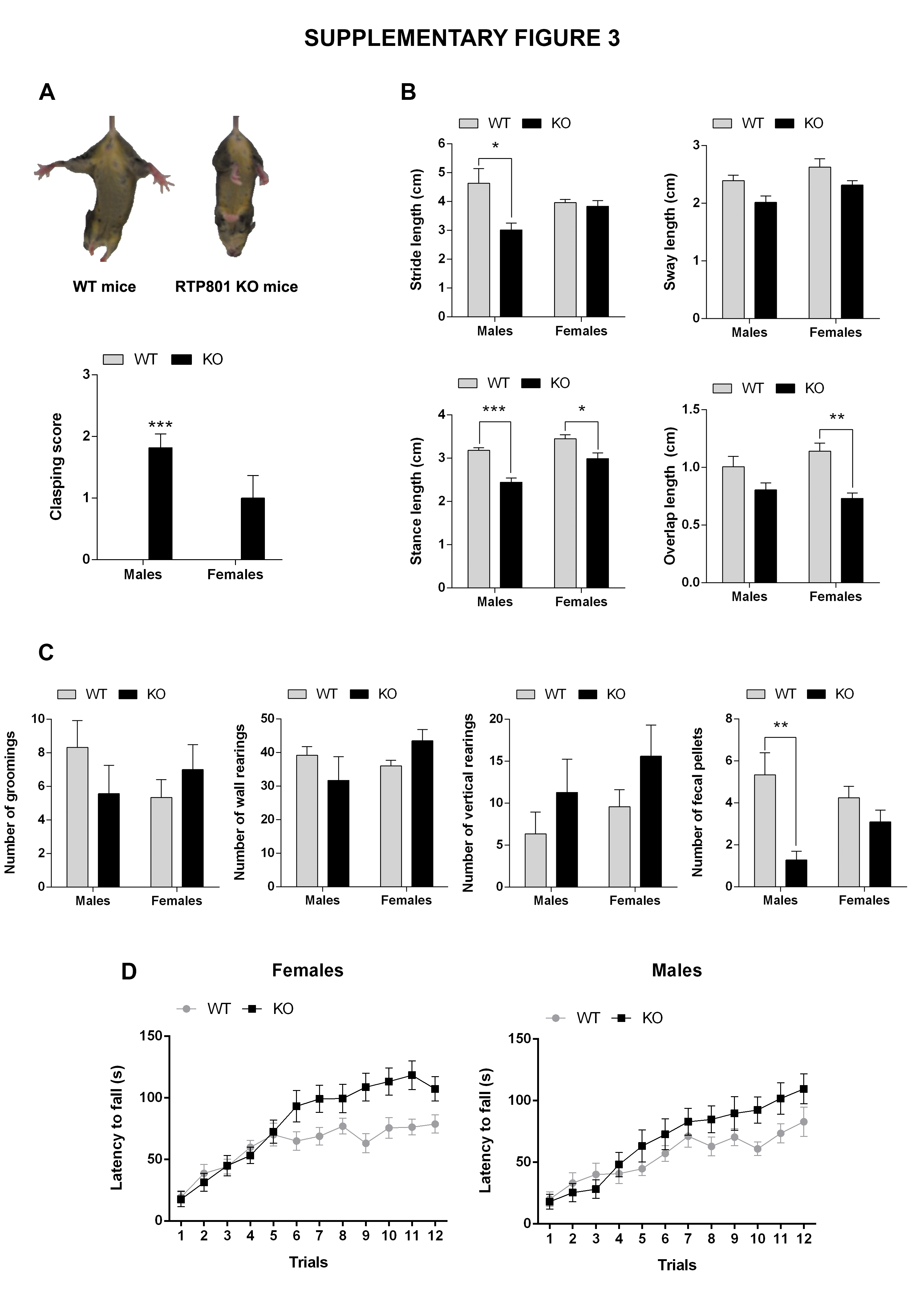

### S4

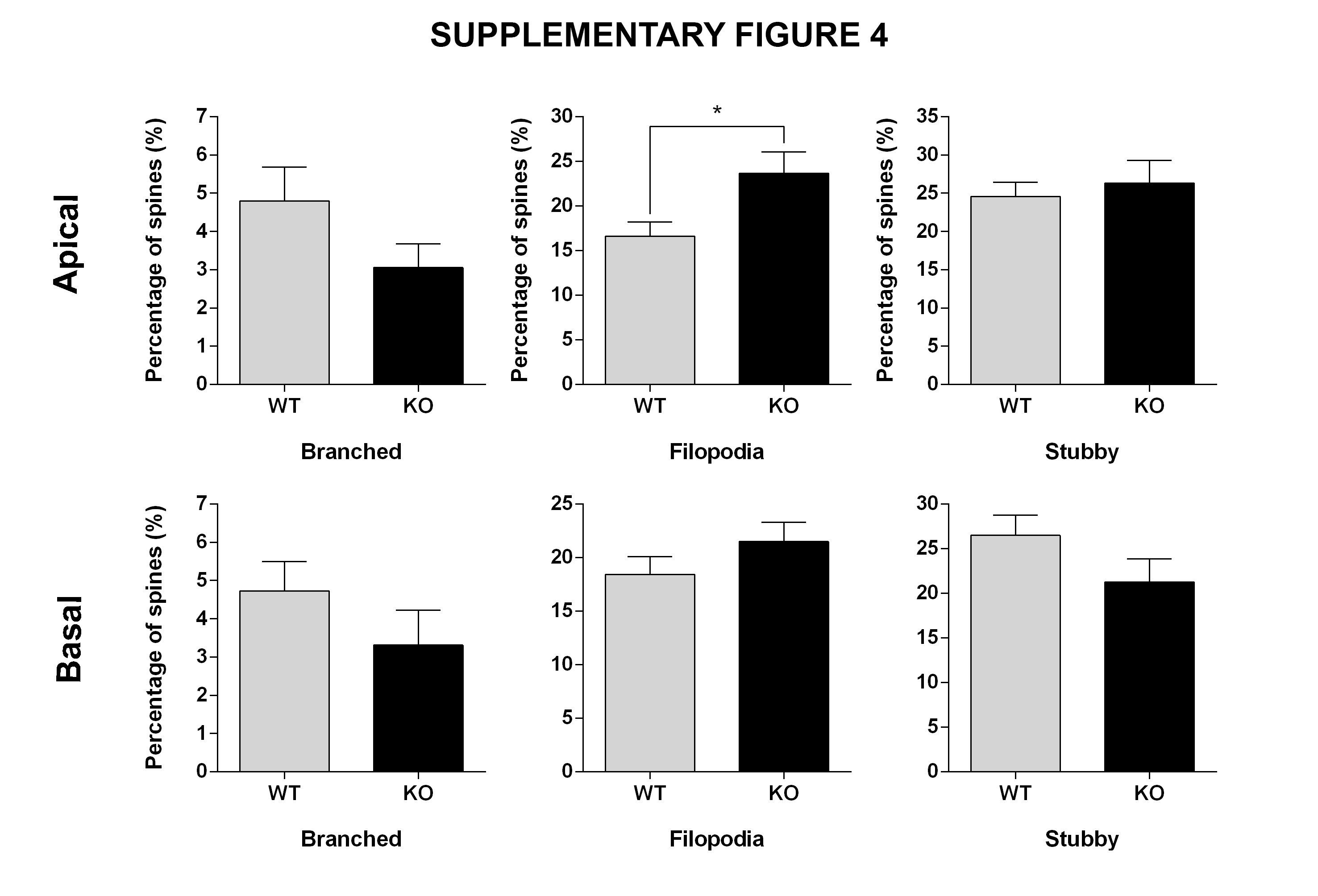

### S5

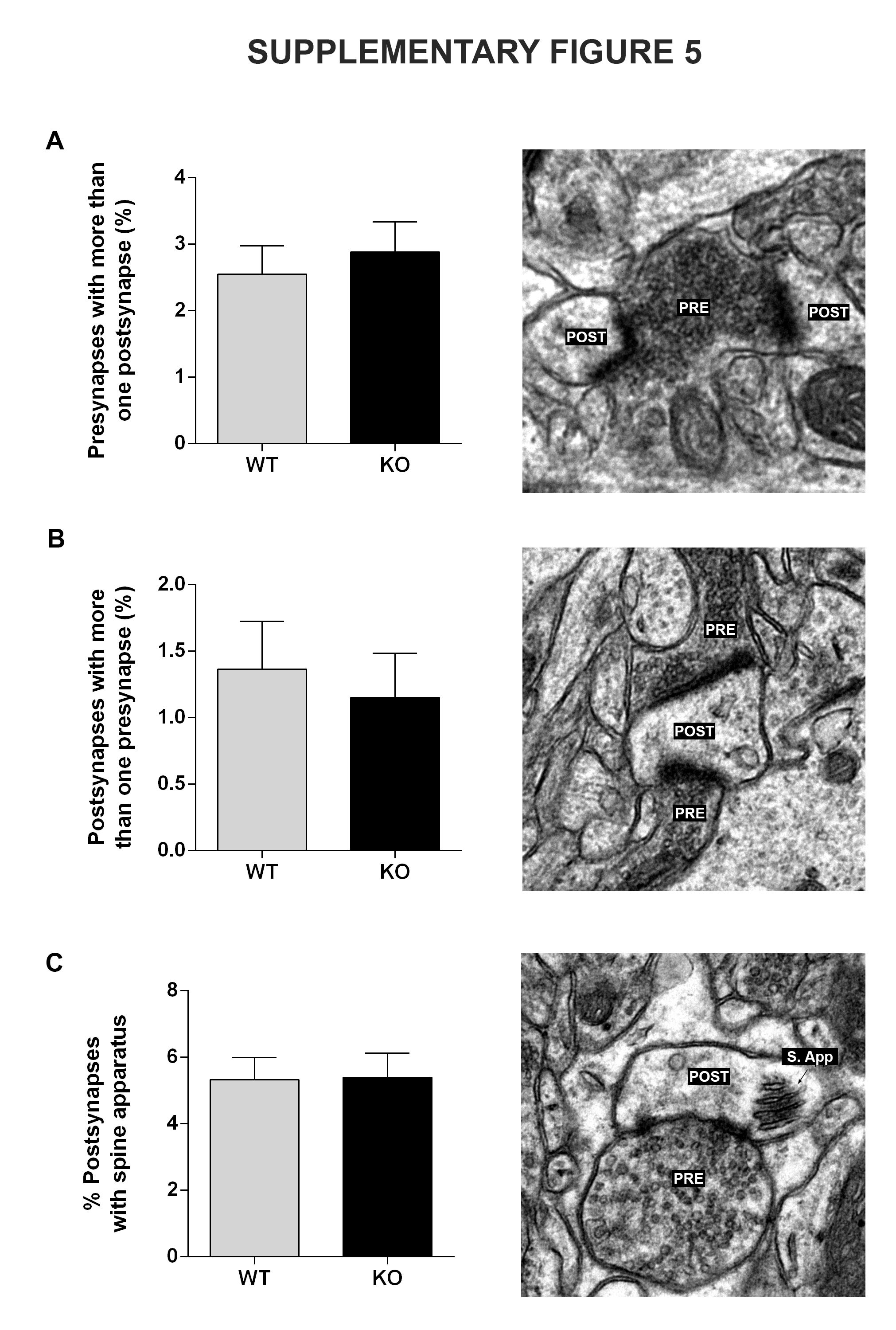
